# One-step induction of human GABAergic neurons promotes presynaptic development & synapse maturation

**DOI:** 10.1101/2025.06.30.662293

**Authors:** Torben W. van Voorst, Maaike A. van Boven, Kevin I. Marinus, Jennifer M. Colón-Mercado, Jule Schretzmeir, Carolin Haag, Ruud F. Toonen, Frank Koopmans, Michael E. Ward, August B. Smit, Ronald E. van Kesteren, Matthijs Verhage, L. Niels Cornelisse

## Abstract

Human induced pluripotent stem cells (iPSCs) present a powerful approach to study human brain physiology and disease, yet robust, pure GABAergic induction has remained difficult. Here we present improved, single-step, transposon-based GABAergic induction with Ascl1/Dlx2, which yields, unlike lentiviral approaches, exclusively GABAergic neurons and was validated across three independent iPSC lines. Co-seeding with Ngn2-induced excitatory neurons created stable networks of predefined excitation/inhibition ratios, with corresponding synapse ratios. Proteomic and electrophysiological characterization at different developmental time points showed that the single-step induced GABAergic neurons gain a proteomic profile that maps to different cortical interneuron subtypes and display typical GABAergic synaptic properties, producing large, synchronous and picrotoxin-sensitive currents. During early development, synaptic strength increased threefold, which was accompanied by an increase in expression of proteins exclusively enriched for presynaptic SYNGO terms. Synaptic strength continued to increase during late development but with only minor proteomic changes. Taken together, transposon-based GABAergic induction yields exclusively mature GABAergic neurons suitable for studying genes involved in synaptic maturation and to build excitation/inhibition networks for disease modelling.

## Introduction

Human induced pluripotent stem cell (iPSC)-derived models provide a promising platform for studying human brain physiology and pathology. iPSCs can be generated from patient-derived tissue or blood to better reflect the human genetics and (patho)physiology compared to commonly employed rodent models.^1–3^ GABAergic dysfunction and excitation/inhibition (E/I) disbalance have been implicated in many neurodevelopmental and neurodegenerative disorders.^4–9^ In recent years, various strategies have been developed for generating iPSC-derived GABAergic neurons, but robust, pure GABAergic induction remains challenging. Directed differentiation, which employs signalling molecules to mimic neurodevelopment.^10–12^, yields both glutamatergic and GABAergic neurons, but these type of inductions often generate inconsistent E/I ratios, tend to yield heterogeneous and immature neuronal populations and are costly and time-consuming.^13^ Rapid differentiation approaches rely on overexpression of key transcription factors, such as Ascl1 and Dlx2 for GABAergic differentiation.^14,15^ and often rely on lentiviral delivery of transgenes. However, such approaches have several limitations: (1) they require multiple vectors when delivering multiple transgenes due to the size cap of ∼8 kb, (2) they pose a risk of disrupting host genes^16^, and (3) they are susceptible to epigenetic silencing during prolonged iPSC culturing ^17–19^ To overcome these challenges, TALEN- and CRISPR-based gene editing.^20^ have been used to target the transgenes to the *AAVS1* safe harbor locus^21,22^ While this ensures stable transgene expression, allows for delivery of multiple genes in a single construct, and minimizes off-target effects, it is time and resource-consuming due to the need for clonal expansion and extensive validation of every iPSC line used in a study.^23^

The PiggyBac transposon system has emerged as a gene-delivery tool that overcomes most of these limitations. Unlike lentiviral systems, transposons enable stable genomic integration of large genetic constructs up to 200 kb.^24^ through a “cut-and-paste” mechanism^25^, which enables delivery of multiple transgenes in a single construct. Additionally, transposon-delivered transgenes are less prone to silencing than lentivirally delivered transgenes^26^ and the increased packing capacity permits the inclusion of epigenetic regulators to further diminish silencing. Compared to safe harbor insertion of transgenes, the delivery of a transposon cassette is a simpler, single-step process applicable to multiple iPSC lines in parallel and allows the use of a polyclonal population for most use cases, eliminating the need for thorough validation, thereby accelerating experimental timelines.

Here, we characterize PiggyBac transposon-based overexpression of *Ascl1* and *Dlx*2 (PB-AD) for GABAergic neuronal induction and benchmark this against established GABAergic induction methods. PB-AD-based induction results in exclusively GABAergic differentiation across different genetic backgrounds, unlike lentiviral approaches which induce both GABAergic and glutamatergic differentiation. We demonstrate that PB-AD neurons integrate into synaptically connected E/I networks when co-cultured with excitatory NGN2 neurons with synaptic projections proportional to cellular ratios. Using proteomics, patch-clamp electrophysiology, and immunocytochemistry we show that PB-AD induction yields neurons with reliable, plastic synapses that synchronously release GABA and strengthen during prolonged culture, with pronounced presynaptic proteomic changes. We propose that this improved GABAergic induction protocol, which reliably generates pure GABAergic neurons, enables the study of human GABAergic functioning in pure GABAergic networks, in single-neuron (autaptic) cultures, and stable E/I-networks.

## Results

### GABAergic differentiation of human iPSCs using PiggyBac based delivery of Ascl1 and Dlx2

Overexpression of the transcription factors Ascl1 and Dlx2 drives GABAergic lineage specification of induced pluripotent stem cells (iPSCs)^15^ Previous approaches employed separate lentivectors to express both genes in an inducible manner. We aimed to develop a simplified GABAergic differentiation protocol using a single PiggyBac transposon construct containing both transcription factors (RRID: Addgene_182307,Figure 1A). This vector contains (1) a doxycycline (dox)-inducible expression cassette for *Ascl1* and *Dlx2* from a bidirectional promoter, avoiding variation in expression levels resulting from position effects in multi-cistronic vectors; (2) reverse tetracycline-controlled transactivator (rtTA), required for dox-inducible expression; (3) the puromycin resistance gene (pac), allowing for antibiotic selection, as well as (4) mTagBFP2^27^, giving transfected cells a fluorescent marker; and (5) Ubiquitous Chromatin Opening Elements (UCOE)^28^ and 5’HS chicken β-globin (cHS4) insulator elements.^29^ Firstly, we tested whether our newly designed construct could induce GABAergic differentiation (Figure 1B). To this end, we co-transfected this PiggyBac vector with a transposase vector into BIONi010-C iPSCs.^30^ After puromycin-selection, all cells expressed mTagBFP2 (Figure 1C), confirming successful transgene integration. Upon dox-induction, cells first clustered together (Figure 1D) and after releasing and reseeding at 3 days post-induction (DPI), cells developed a neuronal morphology, which became more distinct within one week (Figure 1E). At 10 weeks post-induction (WPI), neurons displayed MAP2+ dendrites and VGAT^+^ puncta, indicative of GABAergic synapses, whereas glutamatergic VGLUT1/2^+^ puncta were absent (Figure 1F,G).

**Figure 1.**
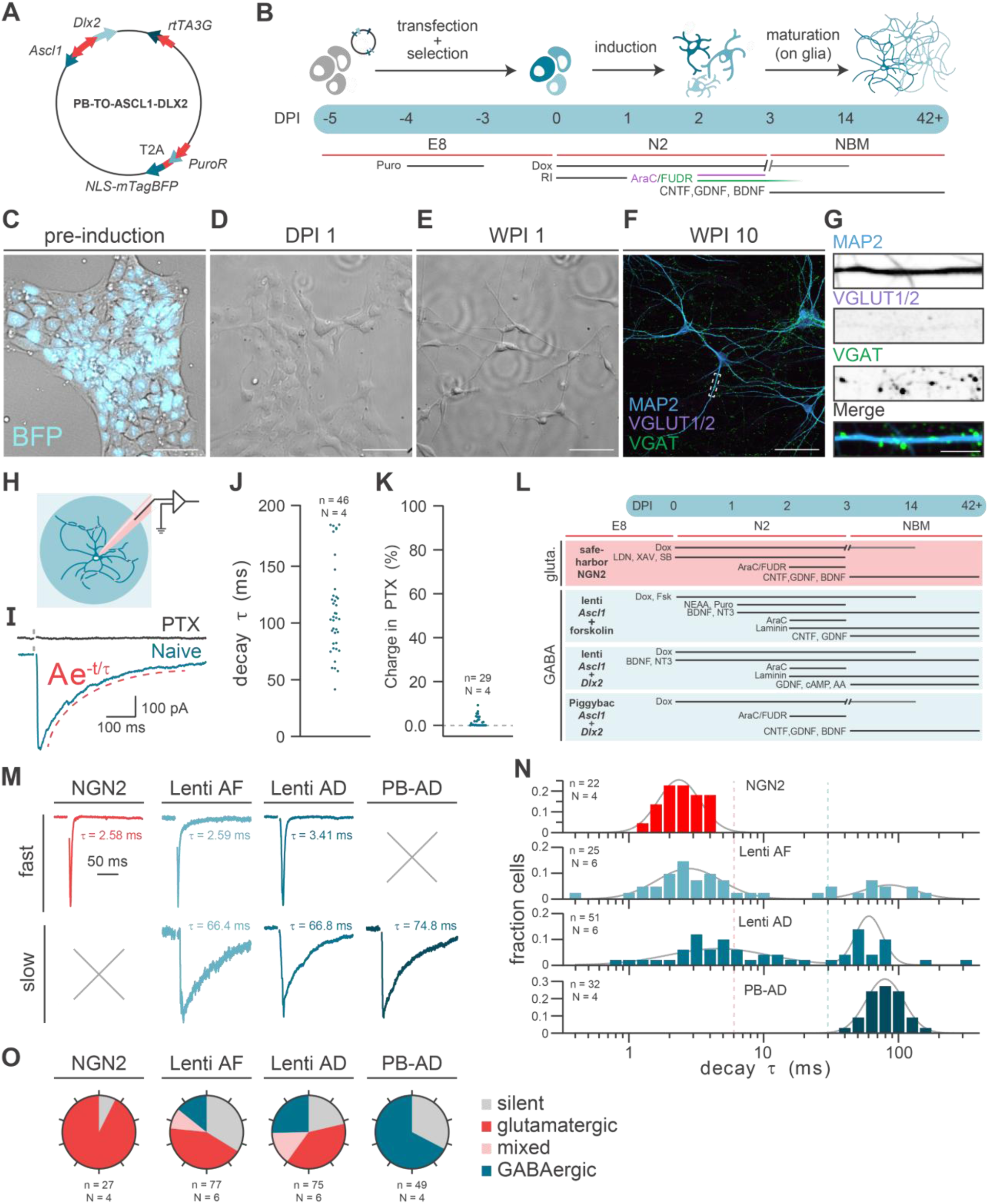
PiggyBac Asc1/Dlx2 efficiently generates pure GABAergic neurons, **A** Map of the all-in-one PiggyBac (PB) vector, expressing murine *Ascl1* and *Dlx2* from a bi-directional TRE-responsive promoter, rtTA from the CAG promoter and *pac*-T2A-NLS-mTagBFP from the EF1α promoter. **B** Overview of the protocol for *Ascl1*/*Dlx2* induction. **C** Before induction, all iPSCs show nuclear BFP signals, demonstrating successful integration of the PB construct **D** Dox-induced expression of *Dlx2* and *Ascl1* triggers neuronal morphology to arise within one day (DPI1) **E** A week after induction (WPI 1), neural morphology is clearly visible **F** After 10 weeks of culture, PB *Ascl1*/*Dlx2* induced neurons show mature neuronal morphology with GABAergic synapses (VGAT+ puncta) and no glutamatergic synapses (VGLUT+ puncta) Scale bar: 100 µm **G.** Zoom-in on the dendrite highlighted in F. scale bar = 1 µm. **H** Autaptic cultures of PB *Ascl1*/*Dlx2* induced neurons were assessed using patch clamp electrophysiology. **I** Evoked postsynaptic currents (PSCs) were fitted using a mono-exponential decay function (red) and their currents were diminished by the application of PTX. **J** Dot-plot of the decay time constants of ePSCs recorded in PB *Ascl1*/*Dlx2* induced neurons. **K** Dot-plot showing the distribution of cells with 0-10% of the naïve charge remaining in PTX **L** Overview of the four induction methods used, PB *Ascl1*/*Dlx2* induction was compared to the lentiviral *Ascl1*/*Dlx2* induction, *Ascl1*+Forskolin induction and *NGN2* glutamatergic induction. **M** Example ePSC traces with either fast or slow decay, as measured in the inductions tested here. **N** Histogram of log-transformed decay time constants (decay τ) measured in neurons induced with the four different induction strategies, together with a ‘sum of two gaussians’ fit (grey line). **O** Pie-charts showing the percentages of neurons with a slow (τ ≥ 30 ms), intermediate (6 > τ < 30 ms) and fast (τ ≤ 6 ms) decay, as well as the percentage of synaptically silent neurons, of neurons generated with the four different induction strategies.

### GABAergic synaptic transmission in PiggyBac Ascl1/Dlx2 neurons

To confirm the GABAergic identity of PiggyBac-induced neurons we performed voltage-clamp recordings in autaptic cultures.^31^ between 6 and 10 WPI (Figure 1H). Rapid depolarization to +30 mV induced action potential evoked postsynaptic currents (ePSCs), which displayed slow decay times (τ) characteristic of GABAergic currents^32^, as determined by mono-exponential fits (geometric mean (GM) = 104.2 ms, geometric SD factor (GM^SDf^) = 1.42; Figure 1I,J). Furthermore, virtually all synaptic signalling was abolished after applying the GABA receptor antagonist picrotoxin (PTX), confirming GABAergic synaptic identity (Figure 1I,K).

### PiggyBac Ascl1/Dlx2 yields a pure GABAergic neuron population in contrast to lentiviral methods

To benchmark the PiggyBac approach, we compared it with two established lentiviral methods for GABAergic induction^14,15^, and an NGN2-based protocol for glutamatergic induction^22,33^ (Figure 1L). Autaptic neurons were generated using all four protocols and ePSC decay time constants were quantified using mono-exponential fits. As expected, NGN2-derived neurons exhibited fast ePSCs decay kinetics typical of glutamatergic transmission (GM = 2.43 ms, GM^SDf^ = 1.39, Figure 1M,N), confirmed pharmacologically and by the presence of VGLUT1/2^+^ puncta (Figure S1). Surprisingly, decay time constants from neurons generated with the lentiviral GABAergic protocols showed a bimodal distribution which could be fitted with a sum of two gaussians. More than half of the cells in these cultures exhibited ePSC kinetics comparable to those in the NGN2 cultures (Lenti AF: GM = 2.82 ms, GM^SDf^ = 0.58, 72.8% of the fit; Lenti AD: GM = 4.62 ms, GM^SDf^ = 0.42, 55.3% of the fit). The remaining cells had more than tenfold larger time constants typical for slow GABAergic ePSCs (Lenti AF: GM = 86.3 ms, GM^SDf^ = 1.56 for 27,2% of the fit; Lenti AD: GM = 60.5 ms, GM^SDf^ = 1.26 for 44.7% of the fit). In contrast, the PB-AD method produced a unimodal population with slowly decaying ePSCs (GM = 78.9 ms, GM^SDf^ = 1.35). Since NGN2 and PB-AD neurons produced decay time constants with unimodal fits, these values were used to determine cut-off thresholds for synaptic identity (glutamatergic: τ ≤ 6 ms; GABAergic: τ ≥ 30 ms; mixed: 6 < τ < 30 ms). This revealed that 68% of PB-AD neurons exhibited GABAergic currents, while the remaining 32% were synaptically silent (Figure 1O). In contrast, the lentiviral approaches generated a substantial population of glutamatergic (Lenti AF: 43%, Lenti AD: 39%) and mixed (Lenti AF: 9% Lenti AD: 15%) identities. The consistent generation of functional GABAergic neurons and the absence of glutamatergic neurotransmission underscore the reliability of the PB-AD protocol compared to previously described methods.

### PiggyBac Ascl1/Dlx2 protocol induces GABAergic neurons across genetic backgrounds

To test for generalizability, we applied the PB-AD protocol to generate neurons from two additional iPSC lines, KOLF2.1J^34^ and GM23973^35^, and characterized their functional properties at 7 and 6 WPI, respectively. In both lines the PB-AD approach produced neurons with GABAergic synaptic projections and slow-decaying synaptic transmission, indicative of GABAergic specification (Figure S2), as was found for the BIONi010-C line. These findings demonstrate the protocol’s applicability across genetic backgrounds.

### PiggyBac Ascl1/Dlx2 neurons integrate into controlled excitatory/inhibitory networks

To test whether PB-AD neurons integrate into E/I networks alongside glutamatergic neurons, we co-cultured 3 DPI PB-AD neurons with 3 DPI glutamatergic NGN2 neurons in networks containing 100%, 30% (as found in the human cortex^36–38^) and 0% inhibitory neurons (Figure 2A-C). Immunocytochemistry at 10 WPI (Figure 2B,C) revealed comparable neuronal cell densities across all conditions, as measured by NeuN staining, indicating similar viability of PB-AD and NGN2 neurons. (Figure 2D). Total dendrite length per field of view increased as the proportion of PB-AD neurons decreased, consistent with the distinct morphological characteristics of GABAergic neurons^39^ (Figure 2E). Pure PB-AD networks contained VGAT^+^ (GABAergic) synaptic puncta, and NGN2 networks contained VGLUT1/2^+^ (glutamatergic) synaptic puncta, with in both cases virtually no puncta of the other type present (Figure 2F,G). In contrast, the mixed (30% PB-AD) networks contained both GABAergic and glutamatergic puncta on the same dendrite proportional to cellular ratios, demonstrating that PB-AD neurons are viable components for assembling E/I networks when mixed with NGN2 neurons.

**Figure 2.**
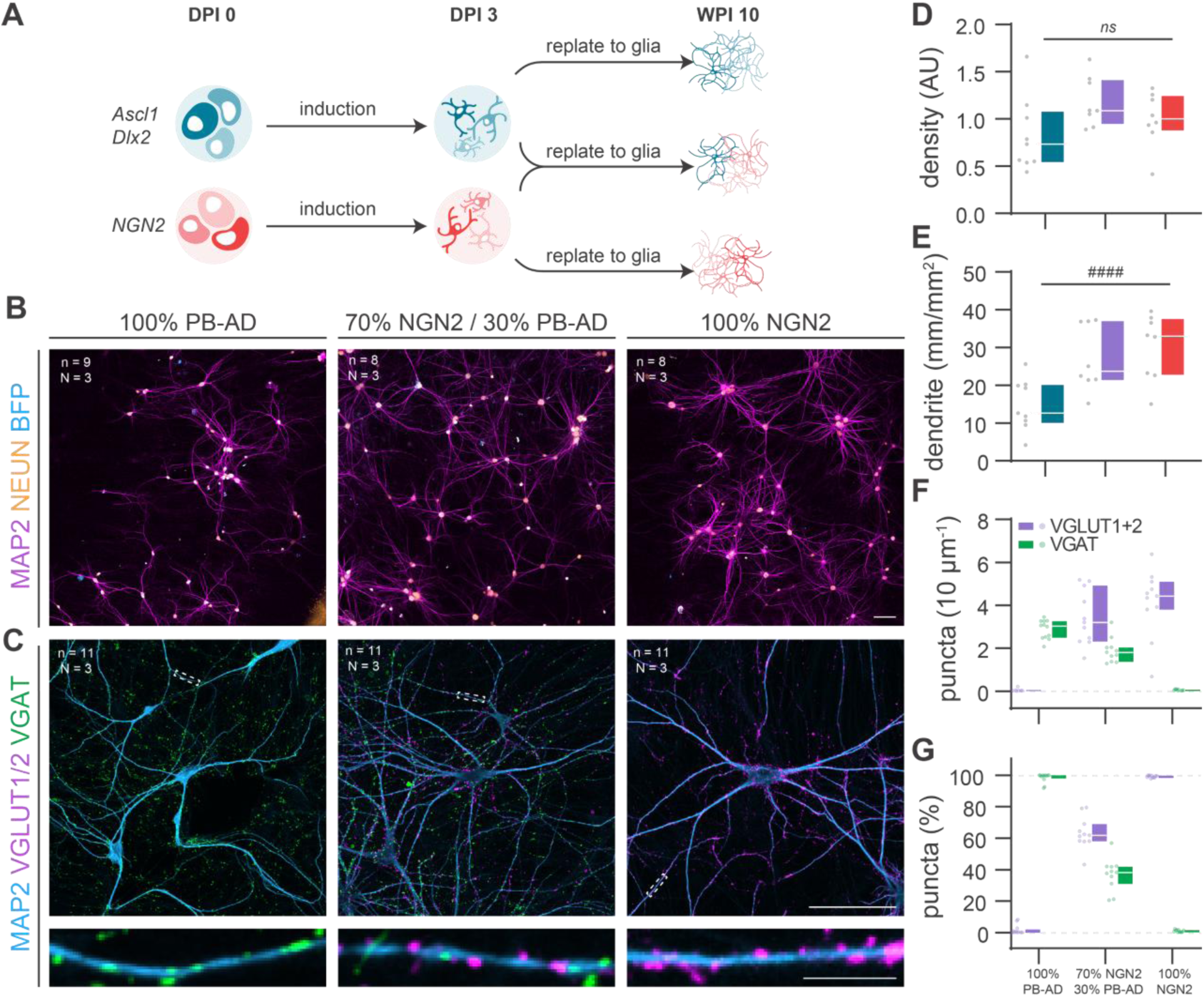
PiggyBac Ascl1/Dlx2 neurons integrate into well-defined excitatory/inhibitory networks upon co-culturing with NGN2 induced neurons. **A** Schematic of co-culturing protocol in which PiggyBac (PB) Ascl1/Dlx2 (AD) neurons were co-cultured with glutamatergic NGN2 induced neurons to generate excitatory/inhibitory (E/I) co-cultures **B** Representative images of E/I co-cultures at different ratios in which dendrites and neuronal nuclei are visualised using MAP2 and NeuN staining, respectively, and PB-AD neurons visualised by nuclear mTagBFP signals. The E/I-ratio is indicated above each image. Scale bar = 100 µm. **C** Representative images of E/I co-cultures at the same ratios as depicted in (B), stained for MAP2 as dendritic marker and VGAT and VGLUT as markers for GABAergic and glutamatergic synapses, respectively. Scae bars = 100 µm, scale bar zoom in = 10 µm **D** The network density (number of nuclei/field of view) is stable across E/I co-culture ratios. **E** The total dendrite length per field of view decreases as the percentage of plated PB-AD neurons increases. **F** The density of glutamatergic (VGLUT^+^ puncta) and GABAergic (VGAT^+^) puncta scales with the percentage of NGN2 and PB-AD neurons plated. **G** Synapse counts shown in F expressed as the percentage of all synapses counted.

### PiggyBac Ascl1/Dlx2 neurons acquire a GABAergic molecular identity over time

Next, we assessed differentiation of PB-AD and NGN2 neurons over time using mass spectrometry-based proteomics. PB-AD co-cultures with rat astrocytes were compared to NGN2 co-cultures at 4, 6, 8, and 10 WPI (Figure 3A). We detected 99,154 human peptide precursors mapping to 7,775 unique proteins across 60 samples that passed quality control. Variability within groups was low-to-moderate with coefficients of variation between 14.6% and 32.2%, reflecting consistent detection across replicates (Figure S3A-B). To remove rat astrocytic signal, subsequent analyses focused on neuronally enriched proteins (1690 proteins), which show >70% neuron-specific expression according to single-cell transcriptomics data of the human cortex (Jostad). Principal component analysis (PCA) on the neuronally-enriched protein abundances (Figure 3B, S3C) revealed clear separation between PB-AD and NGN2 cultures along the first two components (PC1: 71.5% and PC2: 6.1%), as well as the separation of different time points within each culture. More specifically, time points from both PB-AD and NGN2 cultures showed clear separation between 4, 6, and 8 WPI, while cultures at 8 and 10 WPI clustered more closely together, suggesting that proteomic maturation plateaus after 8 weeks.

**Figure 3.**
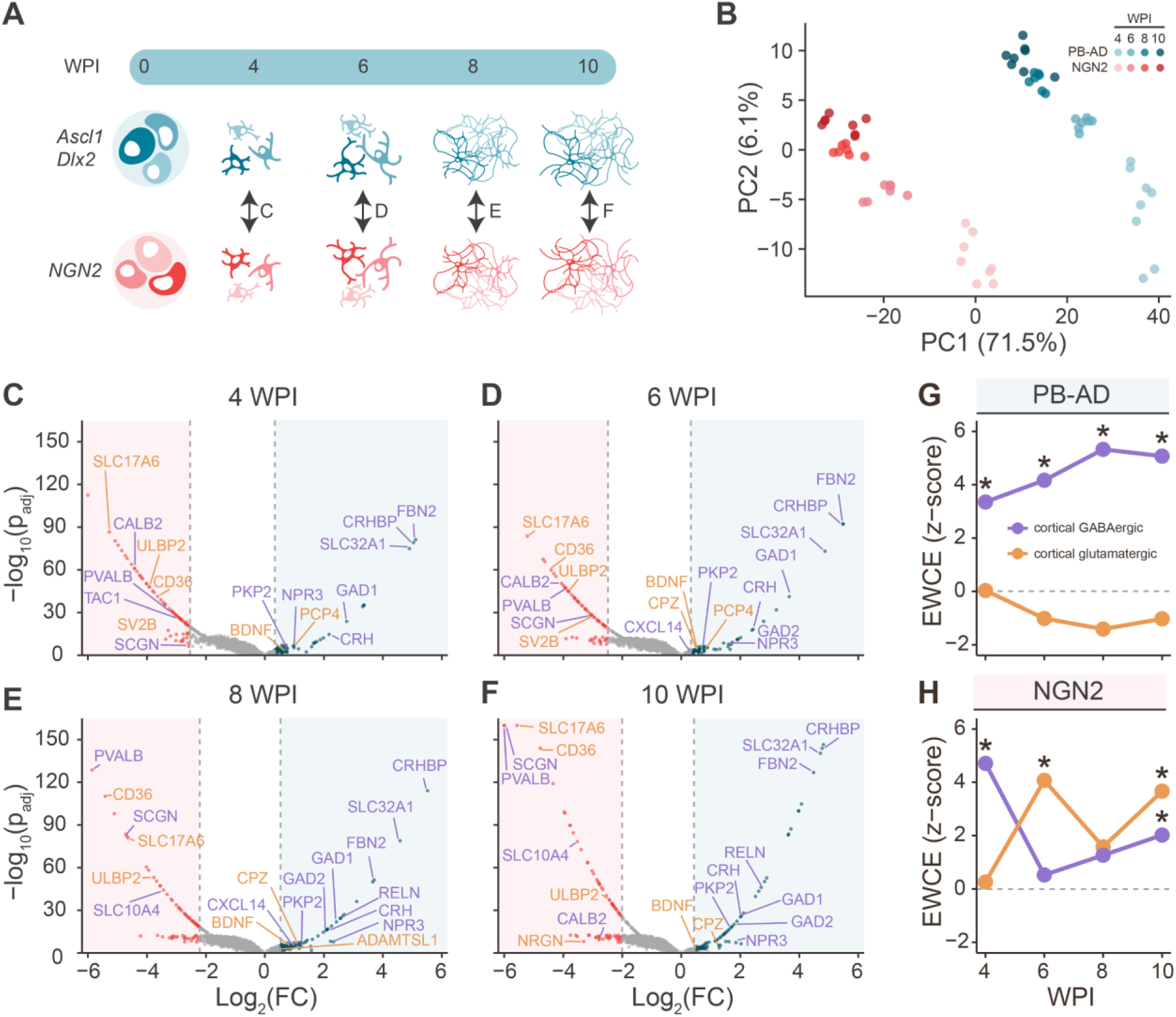
PiggyBac Ascl1/Dlx2 neurons have a cortical GABAergic proteomic profile. **A**. Schematic overview of the proteomics analysis in which PiggyBac (PB) Ascl1/Dlx2 (AD) rat astrocyte co-cultures were compared to NGN2 co-cultures per timepoint. Double-sided arrows indicate the contrasts used for analysis. **B**. PCA analysis on neuronal protein abundances reveals separation of culture types as well as time points along the first two components (PC1: 61.7% and PC2: 8.0%). **C-F**. Volcano plots showing regulation of neuronal proteins in PB-AD cultures and in NGN2 cultures at 4 WPI (**C**), 6 WPI (**D**), 8 WPI (**E**) and 10 WPI (**F**). The top 5% significantly upregulated neuronal proteins in PB-AD are shown as blue dots and those in NGN2 as red dots. Proteins are considered significantly regulated if their adjusted p-value is < 0.05 (significance threshold overlaps with x-axis at this scale and is not shown for clarity of the figure) and their log^2^ (FC) lies above the 95th or below the 5th percentile (vertical dashed lines). Label colours indicate the predicted cell types based on cell type specificity scores (CTSS): regulated GABAergic (purple) and glutamatergic proteins (orange). Cell type specificity was defined using the following CTSS cutoffs: neuronal proteins (neuronal CTSS>70%), GABAergic proteins (GABAergic CTSS > 80% and glutamatergic CTSS < 10%), glutamatergic proteins (glutamatergic CTSS > 80% and GABAergic CTSS < 10%). CTSS can be found in Figure S6. **G**. EWCE analysis of top 5% upregulated neuronal proteins in PB-AD cultures reveals specific enrichment for GABAergic neurons, which increased up to 8 WPI. **H**. EWCE analysis of the top 5% upregulated neuronal proteins in NGN2 cultures reveals significant enrichment for both glutamatergic and GABAergic neurons depending on the timepoint.

To assess lineage specification of PB-AD and NGN2 inductions, we annotated cell types based on differentially expressed proteins using cell type specificity scores (CTSS), computed using single-cell RNA expression data from human cortex^40^ (Figure S4A). Across all timepoints, canonical GABAergic proteins, including SLC32A1 (VGAT), GAD1, GAD2, were enriched in PB-AD cultures. In addition, other predicted GABAergic proteins, such as CRHBP, FBN2, CRH, and RELN, were enriched in PB-AD neurons. In contrast, only a few predicted glutamatergic proteins (e.g., BDNF, PCP4, CPZ, and ADAMTSL1) were moderately enriched in PB-AD cultures. The glutamatergic marker SLC17A6 (VGLUT2) was enriched in NGN2 cultures, while SLC17A7 (VGLUT1) was not detected (Figure 3C-F). Surprisingly, NGN2 neurons also showed strong enrichment of several proteins typically associated with GABAergic neurons at all timepoints, including SCGN, CALB2, and PVALB. Immunocytochemistry confirmed PVALB expression in the majority of NGN2 neurons (Figure S4B).

To quantitatively assess cell type specification, we performed expression-weighted cell type enrichment (EWCE) analysis^41^ on the top 5% upregulated neuronal proteins for each induction for each timepoint. At all timepoints, PB-AD-specific neuronal proteins showed exclusive and significant enrichment for GABAergic neurons, with enrichment levels increasing until 8 WPI (Figure 3G). GABAergic subclass analysis revealed strongest enrichment for VIP^+^ interneurons from 6-10 WPI (Figure S4C). In contrast, NGN2-specific proteins showed significant enrichment for glutamatergic neurons only at 6 and 10 WPI, while also displaying unexpected GABAergic enrichment, reaching significance at 4 and 10 WPI (Figure 3H). Subclass enrichment analysis revealed that this unexpected GABAergic enrichment was most likely accounted for by PVALB^+^ and VIP^+^ interneuron-like profiles (S4C, S4D). Taken together, PB-AD neurons acquire a stable GABAergic identity over time, characterized by consistent expression of canonical and subtype-specific GABAergic proteins. By contrast, NGN2-induced neurons exhibit a mixed molecular identity, with evidence for partial GABAergic specification in addition to expected glutamatergic features.

### Gradual morphological development of PiggyBac Ascl1/Dlx2 neurons

To examine morphological maturation of PB-AD neurons, we performed immunostaining for MAP2, VGAT and VGLUT1/2 in autaptic cultures (Figure S5A,B). Similar to networks, PB-AD neurons produced only VGAT^+^ puncta at all timepoints (Figure S5A,B, Figure 4A-C). A progressive increase in dendrite length was observed during maturation (Figure 4B), accompanied by an increase in the total number of VGAT^+^ puncta per neuron and puncta densities per dendritic length (Figure 4C,D). Additionally, the intensities of VGAT puncta increased over time (Figure 4E), suggesting increased synaptic vesicles per synaptic terminal. Sholl analysis revealed a rise in the number of arborizations during maturation (Figure S5C,D,I). Interestingly, growth of dendritic length and complexity plateaued by 6 WPI, whereas synaptic projections continued to develop until 8 and 10 WPI. Together, these results demonstrate that PB-AD neurons develop complex neuronal morphologies accompanied by an increase in presynaptic GABAergic terminals during maturation.

**Figure 4.**
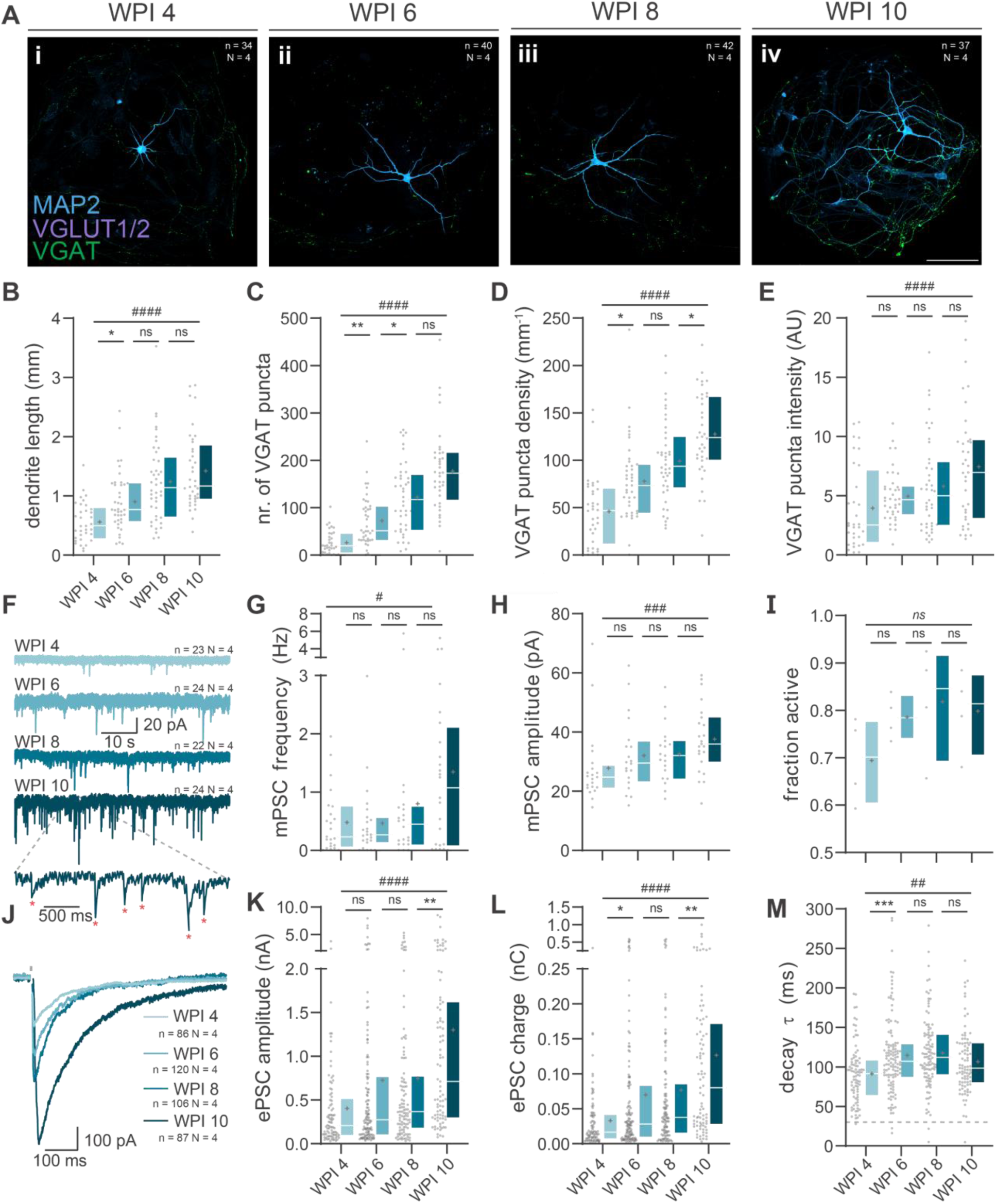
Morphological and functional development of PiggyBac Ascl1/Dlx2 induced neurons. **A** Typical examples of PB-AD autaptic neurons at WPI 4 (i), 6 (ii), 8 (iii) and 10 (iv). Scale bar: 100 µm **B** Total dendrite length quantified. **C** number of GABAergic synapses per neuron. **D** Synaptic density. **E** Average VGAT puncta intensity in arbitrary units (AU). Scale bars: 100 µm. Boxplot whiskers extend the entire data range. **F** Representative traces of mPSCs recorded from PiggyBac (PB) *Ascl1*/*Dlx2* (*AD*) neurons (PB-AD) neurons at 4, 6, 8 and 10 weeks post induction (WPI). **G** mPSC frequency and **(H**) amplitude of PB-AD neurons at 4, 6, 8 and 10 WPI. **I** Boxplots of the average fractions of synaptically active cells at 4, 6, 8 and 10 WPI. Markers represent averages of individual inductions. **J** Representative ePSC traces of at 4, 6, 8 and 10 WPI. **K** Amplitude, charge (**L)** and decay time (**M**) constant of ePSCs recorded from PB-AD neurons at 4, 6, 8 and 10 WPI.

### Synaptic strengthening during maturation of PiggyBac Ascl1/Dlx2 neurons

Next, we assessed functional maturation in autaptic neurons with voltage-clamp electrophysiology. Firstly, action-potential independent spontaneous release was recorded. Both miniature postsynaptic current (mPSC) frequency and amplitude increased during maturation (Figure 4F-H). Next, neurons were depolarized to induce ePSCs. Across all time points, 70-80% of neurons were synaptically active, producing a charge of at least 1.5 pC within 70 ms after stimulation (Figure 4I). In active cells, both the amplitude and charge of ePSCs increased progressively between 4 and 10 WPI (Figure 4J-L), with the most pronounced change occurring from 8 to 10 WPI. Decay constants remained consistently above 30 ms – characteristic of GABAergic currents (Figure 4M). Interestingly, decay kinetics subtly slowed after 4 WPI, suggesting potential changes in postsynaptic receptor subunits.^42,43^

To evaluate the impact of maturation on short-term synaptic plasticity, we applied paired-pulse stimulation at 50 ms, 200 ms, and 1000 ms intervals (Figure S5E-H). This revealed modest synaptic depression which was stable throughout maturation. Only at a 1000 ms interval, synaptic depression increased as neurons matured. Importantly, maturation increased the size of the readily releasable pool (RRP), as determined by back-extrapolation of the charge evoked by 20 Hz stimulus trains, with the biggest change occurring from 8 to 10 WPI (Figure S5I-K). Taken together, these findings demonstrate that PB-AD GABAergic neurons develop functional, plastic synapses that strengthen during maturation.

### Functional maturation correlates with upregulation of proteins in presynaptic gene sets

To investigate the molecular basis of synaptic maturation, we looked at the differential expression of SYNGO-annotated synaptic proteins^44^ between different timepoints of development (4 to 6 WPI, 6 to 8 WPI, and 8 to 10 WPI) (Figure 5A). Of all 1605 SYNGO-annotated synaptic genes, 929 unique proteins were detected in PB-AD cultures, of which 478 (51%) were differentially expressed in at least one time contrast (Figure 5B, D-F). In NGN2 neurons, 967 unique SYNGO-annotated proteins were identified of which 644 (67%) were differentially expressed in at least one time contrast (Figure 5C, Figure S6A-C). In both cultures, SYNGO-annotated proteins were abundantly regulated between 4 and 8 WPI, while only few proteins showed altered expression between 8 and 10 WPI (Figures 5B-F, Supplementary table 1). This indicates that proteomic synaptic maturation plateaus after 8 WPI, consistent with what was observed for neuronal-enriched protein expression.

**Figure 5.**
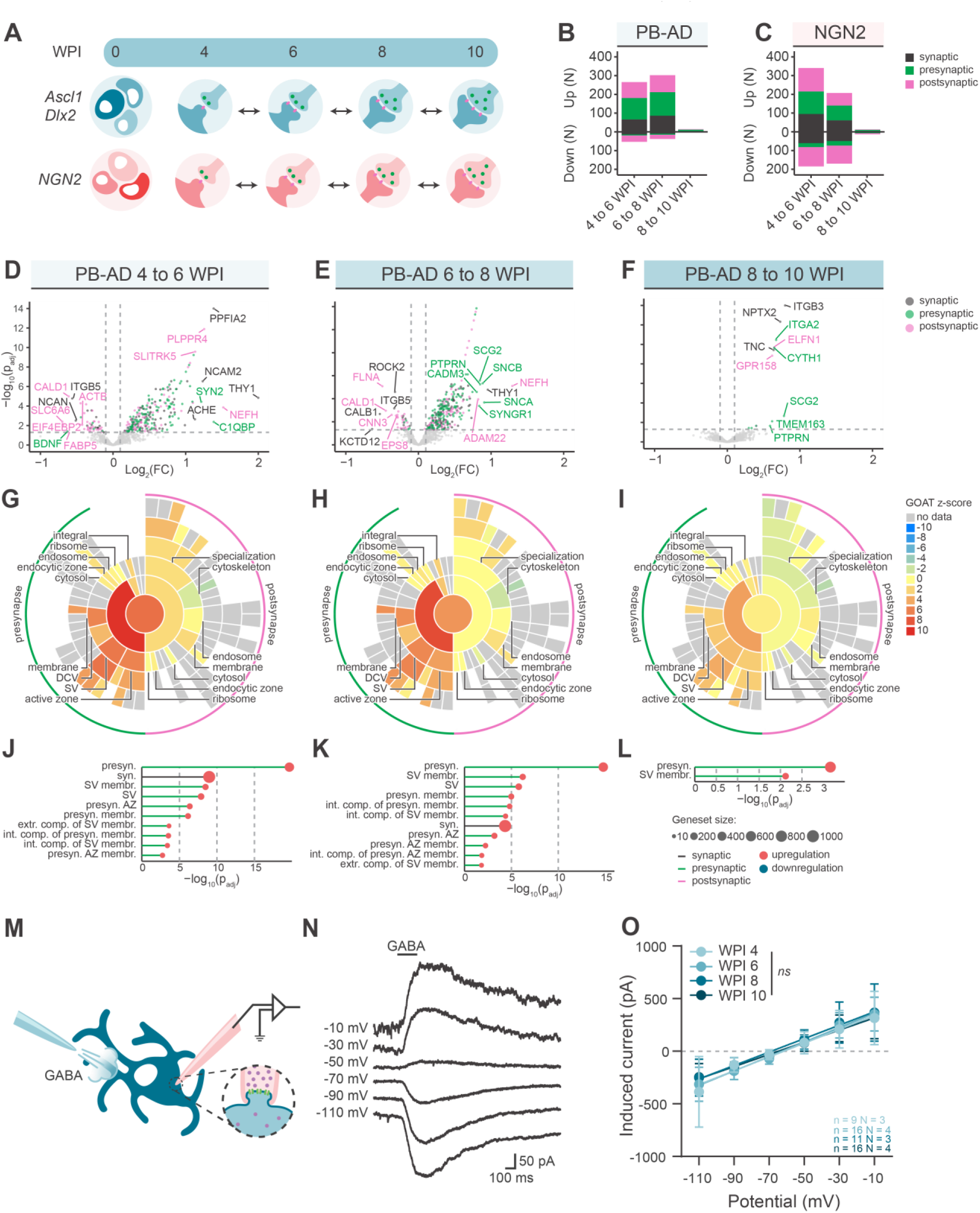
Pre- and postsynaptic maturation of PiggyBac Ascl1/Dlx2 neurons. **A**. Schematic overview of the proteomics analysis with double-sided arrows indicating the contrasts used for analysis. **B-C**. Stacked bar plots showing the number of up- and downregulated SYNGO annotated proteins over time, distinguishing synaptic (black), exclusively presynaptic (green) and exclusively postsynaptic (magenta) proteins for PB-AD (**B**) and NGN2 (**C**) neurons. Classifications were assigned based on SynGO annotations, synaptic classification contains both pansynaptic and ambiguously localized synaptic proteins. Regulated proteins have a log^2^ (FC) > 0.1, or < −0.1, and adjusted p-value < 0.05. **D-F.** Volcano plots showing regulation of synaptic proteins between 6 and 4 (**D**), 8 and 6 (**E**), and 10 and 8 WPI (**F**). Only SYNGO-annotated proteins are plotted and colour coding refer to significantly regulated synaptic (black) exclusively presynaptic (green) or postsynaptic (pink) annotations. Proteins were regarded significantly regulated when adjusted p-value < 0.05 (dashed horizontal line) and log^2^ (FC) > 0.1 or < −0.1 (dashed vertical lines). Top 5% proteins regulated proteins were labelled based on 97.5^th^ and 2.5^th^ percentiles marking genes that drive subsequently detected enrichments. **G-I**. SYNGO plots showing enrichment scores (z-scores) derived from gene-set enrichment analysis with GOAT on SYNGO cellular compartment gene sets, using differential expression output of 6 vs 4 (G), 8 vs 6 (H) and 10 vs 8 WPI (I),. Negative z-scores reflect enrichments primarily driven by downregulated proteins and vice versa for positive z-scores. **J-L.** Lollipop charts showing significant ontologies with their corresponding adjusted p-values and the number of genes part of the gene set as analysed by GOAT for 6 vs 4 (J), 8 vs 6 (K) and 8 vs 10 WPI (L) **M.** Schematic overview of the GABA puff experiment, in which perforated-patch was performed on PB-AD neurons in E/I-networks and GABA was applied to measure the postsynaptic current it produced. **N.** Typical example of the postsynaptic current produced by GABA puff at different holding voltages. **O.** Linear fit of PSC amplitudes as measured in response to GABA puff at 4, 6, 8 and 10 WPI. Error bars represent the SD.

To examine specifically how pre- and postsynapses develop over time, we used GOAT^45^ to determine which SYNGO gene sets were enriched for developmentally regulated proteins, where the contribution of each protein was weighted by its log^2^(fold change).^45^ Strongest enrichment was observed between 4-6 WPI and 6-8 WPI, which occurred exclusively in presynaptic gene sets, including *synaptic vesicle* (*SV*), *presynaptic active zone (AZ)*, and *presynaptic membrane* (Figure 5G-L). Between 8-10 WPI only *SV membrane* proteins and its mother term *presynaptic* proteins were enriched. In contrast, no postsynaptic gene sets were significantly enriched in PB-AD cultures, across all time contrasts. To exclude that these results were biased by a few proteins with large effect sizes, additional SYNGO gene set overrepresentation analysis was performed based on the number of upregulated proteins per gene set. This confirmed the exclusive enrichment of presynaptic terms in PB-AD cultures (Figure S6J–L). Upregulated proteins in NGN2 cultures over time similarly showed a preferential enrichment of presynaptic gene sets using GOAT and SYNGO overrepresentation analyses, with no enrichment for postsynaptic gene sets between 6-10 WPI. Thus, synaptic strengthening correlates with changes specifically in presynaptic protein levels, while postsynaptic protein levels remain relatively stable over time.

In order to test whether our findings on asymmetrical pre- and postsynaptic maturation translate to a more physiological model, we assessed postsynaptic responses to externally applied GABA in E/I networks, as excitation might alter maturation profiles.^46^ (Figure 5M). Furthermore, we used gramicidin-perforated patch to allow measurement of postsynaptic responses without influencing the reversal potential for the GABA-receptor^47^ GABA puffs were applied at varying holding voltages (Figure 5N). The slope of the I/V curve voltages was unaltered by maturation (p = 0.305), indicating that the GABAergic postsynaptic response does not change during maturation under these conditions. Thus, increased synaptic strength during synaptic maturation correlates with presynaptic proteomic changes, but not with postsynaptic development also in E/I-networks.

### PiggyBac Ascl1/Dlx2 specific synaptic proteins map to cortical interneuron subtypes

Finally, we investigated whether changes in PB-AD and NGN2 SYNGO-annotated proteomes between 4 and 10 WPI are associated with cortical GABAergic and glutamatergic cell types.^40^ (Figure 6A). PB-AD cultures showed upregulation of predicted GABAergic and medial ganglionic eminence (MGE) and caudal ganglionic eminence (CGE)-specific proteins (Figure 6B). NGN2 cultures showed upregulation of predicted glutamatergic proteins (Figure 6C). Of note, PB-AD and NGN2 cultures showed upregulation of both GABAergic and glutamatergic receptor subunits (Figure S7A,B), although only input of one of the neurotransmitters was present per culture. Canonical subtype markers for both MGE- and CGE-derived subtypes were also detected in PB-AD neurons (S7C,D).

**Figure 6.**
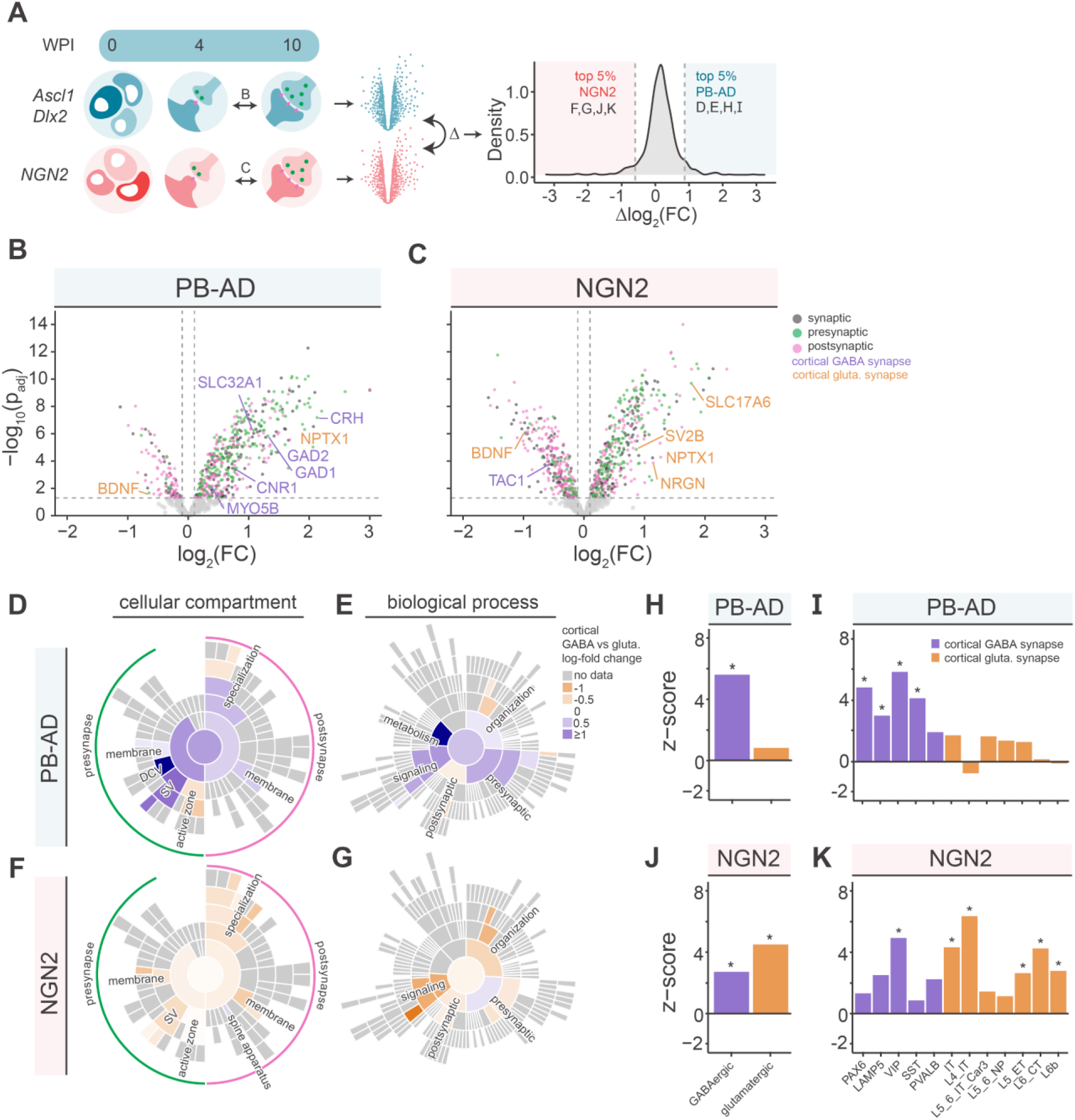
Specialization of PiggyBac Ascl1/Dlx2 and NGN2 neurons into cortical subtypes. **A.** Schematic overview of the proteomics analysis pipeline. The differential expression of synaptic proteins over time (10 vs 4 WPI) was first quantified for PB-AD and NGN2 culture separately. The difference in expression over time between PB-AD and NGN2 (delta-log^2^ (FC)) was next calculated, enabling identification of proteins with PB-AD- or NGN2-specific temporal profiles. Density plot showing 5^th^ and 95^th^ percentiles (dashed vertical lines) selections of proteins that have strong differential temporal regulation (i.e., high or low delta-log^2^ (FC)) in PB-AD compared to NGN2, and vice versa. **B-C**. Volcano plots showing regulation of synaptic proteins between 10 and 4 WPI. Proteins are considered regulated when the log^2^ (FC) > 0.1 or < −0.1 (dashed vertical lines) and adjusted p-value < 0.05 (dashed horizontal line). Only SYNGO-annotated proteins are plotted, and dot colors refer to significantly regulated synaptic (black) exclusively presynaptic (green) or postsynaptic (pink) annotations. Label colours indicate for predicted cortical GABAergic- (purple) or glutamatergic (orange) regulated proteins. **D-G** SYNGO sunburst plots showing the GABAergic and glutamatergic specification of PB-AD and NGN cultures. The SYNGO sunburst inputs are the proteins that are part of the 5^th^ (**F-G**) and 95^th^ percentiles (**D-E**) from the delta-log^2^ (FC) distribution (**A**), which represent the top 5% exclusively PB-AD and NGN2 regulated proteins, respectively. GABAergic or glutamatergic annotation for these SYNGO terms is estimated based on the differential RNA expression (log^2^ (FC)) between cortical GABAergic and glutamatergic neurons of corresponding genes^40^ Both cellular compartment (**D-F**) and biological process (**E-G**) gene sets were used. **H-K**. EWCE analysis of PB-AD-specific (**H-I**) or NGN2-specific proteins (**J-K**) show enrichment for GABAergic and glutamatergic cell sub (classes), respectively. IT and ET refer to intra- and extratelenchephalic-projecting neurons and NP refers to near-projecting neurons.^40^

To analyze lineage-specific synaptic development in more detail, we next contrasted the developing (WPI 4 to WPI 10) PB-AD and NGN2 SYNGO-annotated proteomes with each other to find proteins that were specifically regulated in each induction. To this end, for each protein we calculated the difference between PB-AD and NGN2 in differential expression between 4-10 WPI (i.e., log^2^ (FC)^AD^-log^2^ (FC)^NGN2^), and selected the 5^th^ and 95^th^ percentile to define induction-specific synaptic proteins (Figure 6A). We then assessed whether these proteins were enriched in either cortical glutamatergic or cortical GABAergic reference expression profiles^40^ Overall, this produced lineage-specific enrichments in most gene sets. PB-AD specific synaptic proteins showed the highest GABAergic enrichment in the gene sets *dense core vesicle* (*DCV*), *SV*, *synaptic signalling*, *metabolism* and *presynaptic process* (Figure 6D-E). Conversely, NGN2-specific synaptic proteins that belonged to *synaptic signalling, SV, organization* and *specialization* showed the highest glutamatergic enrichment (Figure 6F-G).

As synaptic gene expression profiles have been shown to reflect neuronal subtypes^48^, we queried whether synaptic proteins that specifically increase in PB-AD or NGN2 cultures show enrichment for neuronal (sub)types. In EWCE analysis, the PB-AD-specific synaptic proteins were enriched in GABAergic neurons (Figure 6H), specifically VIP^+^, SST^+^, PAX6^+^ and LAMP5^+^ interneurons, but not PVALB^+^ neurons (Figure 6I). NGN2-specific synaptic proteins were enriched in glutamatergic subtypes (IT, L4 IT, L5 ET, L6 CT, and L6b) but also in GABAergic subtypes (Figure 6J), particularly VIP^+^ interneurons (Figure 6K). Together, these data indicate that PB-AD-based induction leads to GABAergic synaptic specialization with molecular profiles resembling CGE- and MGE-derived interneurons, whereas NGN2 induction leads to induction of neurons that also contain typical GABAergic proteins.

## Discussion

### Summary

In this work, we present a single-step, transposon-based GABAergic differentiation method, which uses a single vector to overexpress Ascl1/Dlx2, and we provide a detailed characterization of these GABAergic neurons at different developmental timepoints using mass spectrometry proteomics, immunocytochemistry, and patch-clamp electrophysiology. This method is applicable across different genetic backgrounds and produces exclusively GABAergic neurons with a proteomic profile that maps to different cortical interneuron subtypes, particularly VIP^+^ neurons. Synapses display GABAergic properties such as picrotoxin-sensitive, slowly decaying, evoked postsynaptic currents and short-term depression, similar to primary GABAergic neurons.^49–53^ During development, synapses gradually increase in synaptic strength, which at earlier timepoints correlates with increased expression of presynaptic proteins associated with active zone, synaptic vesicle, and presynaptic membrane gene sets. In contrast, little functional and proteomic changes occur in the postsynapse. Late in development, synaptic strength continues to increase, with only minor proteomic changes. PB-AD neurons can be combined with NGN2 neurons to form networks of predefined E/I ratios with corresponding synapse ratios. Taken together, transposon-based GABAergic induction yields GABAergic neurons with improved synaptic properties that can be used to build balanced excitation/inhibition networks, for instance to model diseases in which E/I disbalance has been proposed as a disease mechanism.

### PiggyBac Ascl1/Dlx2 neuronal induction produces exclusively GABAergic neurons

By employing autaptic neurons we were able to detect synaptic output on a single neuron scale, which allowed us to accurately quantify lineage specification and demonstrate how GABAergic transposon-based induction compares to lentiviral strategies. Previous studies using a lentiviral approach also detected non-GABAergic currents^54^ and reported expression of glutamatergic genes^55,56^, while others focused on measurements in E/I-networks^14,15^ or recorded under conditions that blocked glutamatergic transmission^57,58^, making it difficult to compare with our results. As Ascl1 expression, by itself, induces glutamatergic differentiation^59,60^, the balance of Ascl1 and Dlx2 expression is likely important in steering differentiation towards a GABAergic fate. The design of our transposon-based induction incorporates multiple features to facilitate this and thereby avoids unwanted glutamatergic specification. Firstly, both factors are present in one cassette and transcribed from a bidirectional promotor. Secondly, a shared integration site could prevent position effects.^61^ Lastly, insulator elements may reduce gene silencing, which is observed for lentiviral cargo^17–19,62–65^, and less so for transposon delivered transgenes.^26^

### Proteomic profiles of PiggyBac Ascl1/Dlx2 neurons map to cortical interneuron subtypes

Cell type enrichment analysis revealed that neuronal proteins that are higher expressed in PB-AD cultures are significantly enriched for GABAergic neurons at all timepoints, with specific enrichment for VIP^+^ interneurons from 6 WPI onwards. Zooming in on synaptic proteins that are upregulated specifically in PB-AD cultures between 4-10 WPI, we showed significant enrichment for all GABAergic interneuron subtypes except PVALB^+^, suggesting specification towards both MGE- and CGE-derived neurons. Assessment of the neuronal lineages produced by PB-AD induction would require single-cell sequencing approaches. A previous single-cell RNA-sequencing study of lenti AD neurons predominantly found SST^+^ neurons at 4 WPI, but later time points were not investigated.^66^ Perhaps CGE markers only appear further along the maturation process, or transcriptomics do not accurately reflect the molecular identity of these neurons.^67–69^ To further restrict and direct subtype specification during PB-AD induction, additional patterning with small molecules, which was recently developed for safe harbor-AD induction, could potentially be employed.^70^

Ngn2-induction produced cultures with a mixed glutamatergic and GABAergic molecular identity, including canonical GABAergic markers such as SCGN, PVALB, CALB2. Although these markers are sometimes found in certain glutamatergic neurons in the brain^71–76^, it is more likely that their expression is a result of nonspecific differentiation. Nevertheless, virtually no VGAT puncta were identified and only glutamatergic ePSCs were recorded, consistent with previous reports.^21,77–82^

### Enhanced expression of presynaptic proteins during synaptic maturation

Our multilevel analysis of GABAergic cultures at different time points between 4 and 10 WPI, combining patch-clamp electrophysiology, immunocytochemistry, and mass spec proteomics, revealed the following insights about synapse maturation. Firstly, synaptic strength increased approximately 3-fold during this period. A comparable increase in mPSC frequency and RRP suggests that this is due to an increase in the strength and/or number of presynaptic projections, which is further supported by VGAT immunocytochemistry. This developmental trajectory is similar to what has been described for directed-differentiation and safe harbor AD GABAergic autaptic neurons.^33,79^

Secondly, we observed a stark difference between pre- and postsynaptic molecular differentiation. Strong upregulation of *AZ-, SV-* and *presynaptic membrane* proteins occurred, while expression changes in proteins in postsynaptic gene sets were minimal. Asymmetric proteomic development of the pre- and postsynapse was also observed in NGN2 neurons and has been described for iPSC-derived neurons obtained through directed differentiation.^83,84^ In PB-AD neurons, current responses to GABA puffs remained constant over time, indicating that functional postsynapses already exist at 4 WPI and the expression of GABA receptors remains unaltered at subsequent weeks. This fits with the notion that expression of postsynaptic neurotransmitter receptors precedes presynaptic innervation.^85–89^ Release from presynaptic projections subsequently induces clustering of postsynaptic molecules, initiating postsynaptic maturation.^85,90–93^ Indeed, postsynaptic receptor clustering was previously predicted *in silico* to increase mPSC amplitude 1.5-fold, similar to what we found *in vitro*^94^.

Thirdly, although proteomic changes plateau after 8 WPI, both synapse number and synaptic strength continue to increase. Possibly, the *AZ, SV* and *presynaptic membrane* proteins that are upregulated until 8 WPI are sufficient to support synapse formation thereafter. Structural rearrangement and posttranslational modification of synaptic proteins, mediated by e.g. ERK, PKC, PKA, CaMKII and/or RhoA/ROCK signalling^95–109^, could enhance synaptic strength without proteomic changes. Such processes modulate many pre- and postsynaptic proteins, including gephyrin^98–100,107,110^, GABA-receptors^101–105,108–110^, synapsins^96,106^ and microtubules and their associated proteins.^110–115^ Additionally, synaptic strength could be regulated by (trans-synaptic) alignment of functionally related proteins in nanodomains and nanocolumns, enhancing their cooperative interactions.^94,116–122^ Of the few proteins that were upregulated – or for the first time detected – in PB-AD neurons at 10 WPI (Supplementary table 1), we found several cell adhesion molecules (CAMs) (CNTNAP4, SLITRK1, CDH1, EPHA7, ITGB3, NPTX2, ITGA2, ELFN1, IGSF8 and CNTNAP1).^3,123–131^, extra cellular matrix (ECM) proteins (TNC and RELN)^132,133^, and proteins involved in receptor clustering (NPTX2, SLITRK1, PRRT1, ELFN1)^134–137^ Some of which are known to strengthen GABAergic transmission.^123,126^ To summarize, during early development, synaptic maturation and strengthening in iPSC-derived neurons coincides with increased expression of presynaptic proteins compared to their postsynaptic counterparts. At later stages, further strengthening may result from alignment and reorganization of the pre- and postsynaptic compartments.

### A versatile in vitro model for studying the role of inhibition in brain disease

Dysfunction of inhibitory neurons and E/I disbalance are common aspect of neurodevelopmental and neurodegenerative disorders.^5,138–143^ Given the pure GABAergic nature of cultures produced with the PB-AD protocol, neurons generated with this method provide a versatile model to study the role of inhibition and E/I disbalance in these disorders using a human, and patient-own, genetic background. Compared to a safe harbor approach, our method is directly applicable to patient derived iPSCs, which makes it time and cost efficient, and prevents the introduction of clonal gene editing effects. The reliable production of GABAergic autaptic cultures allows for a detailed assessment of interneuron synaptic properties, such as priming, fusion and synaptic plasticity in patient cell lines.^144–146^ Application to disease-lines will be important steps to substantiate the full potential of this method. To this end, PB-AD neurons can be mixed with NGN2 neurons at defined ratios, enabling accurate modelling of E/I networks at any ratio, including those found in the human brain.^36–38^

## Methods

Extended methods are available in the supplementary material.

### IPSC culture and delivery of transgenes

IPSC’s were maintained on Matrigel- or Geltrex coated-dishes in Essential 8 medium. For viral induction, viral particles encoding rTTa, Ascl1, and Dlx2 were added to IPSCs and antibiotic selection was performed 24 hours after transfection. For PiggyBac based strategies, IPSCs were seeded as single cells after which transfection with PiggyBac transposon and transposase plasmids was performed using Lipofectamine STEM. Selection was performed 24 hours post-transfection.

### Neuronal induction

Transgene-selected IPSCs were seeded in N2 medium with doxycycline to induce transgene expression. After 2 days of transgene expression, selection for postmitotic cells was performed with antimitotic agents FUDR or AraC, and cells were collected and replated on rat astrocytes in neurobasal medium supplemented with neurotrophic factors, in which neurons were maintained.

### Electrophysiology

Whole-cell voltage clamp was performed to measure synaptic release. For GABA puffs, perforated patch was performed, maintaining cell intrinsic chloride levels. GABAergic and glutamatergic currents were blocked with picrotoxin and DP-V + DNQX, respectively.

### Immunocytochemistry and morphological analysis

Methanol- or PFA-fixed neurons were stained for MAP2, VGAT, VGLUT1/2, NEUN and Parvalbumin. After confocal microscopy analysis was performed in SynD^147^ and ImageJ.

### Statistics of electrophysiological and morphological data

Statistical testing was performed in Matlab and GraphPad Prism. Developmental trajectories were tested using nonparametric Spearman correlations. For timepoint comparisons, Kruskal-Wallis ANOVAs with Dunn’s correction were performed. GABA puff responses were fitted using a first-order polynomial linear fit and an extra sum-of-squares F-test was employed.

### Mass spectrometry

Cells were scraped, pelleted, and stored at −80°C. Sample were prepared using adapted DNA micro spin column suspension trapping protocol.^148^ Pellets were extracted in SDS buffer and proteins were isolated and trypsin-digested. Peptides were eluted from the column and samples were analysed using liquid chromatography MS.

Spectra were mapped to *in silico* spectral library based on the uniprot human reference proteome peptides using DIA-NN 1.8.1.^149^ Protein abundances were quantified using differential abundance analysis (DAA) of MS-DAP 1.0.5.^150^ Proteins (nearly) exclusive to one group were analysed using MS-DAP differential detection (DD) test. The DEqMS^151^ algorithm was used for statistical analysis and resulting p-values were adjusted using Benjamini–Hochberg False Discovery Rate (FDR) adjustment. DAA and DD results were merged. Gene set enrichment analyses were performed using GOAT online v1.0.2.^152^ and SynGO knowledgebase v1.2.^153^

Cell type-associated proteomic changes were identified by expression weighted cell type enrichment (EWCE) analysis^41^, based on snRNA-sequencing data derived from human cortex.^40^ For cell population classifications we used cell class (e.g., GABAergic neurons) and subclass (e.g. SST^+^ interneurons). Cell type-specificity scores (CTSS) for each gene were used to assign cell type labels.

## Supporting information

Quantifications and statistics on electrophysiological and morphological data

## Acknowledgements

The authors thank Robbert Zalm and Ingrid Saarloos for their help with cloning and plasmid production, Lisa Laan and Desiree Schut for preparation of glia feeder plates, Judith Huijgen and Solange Lopes Cardozo for preparation and maintenance of iPSCs, and Berna Özer and Remco Klaassen for performing mass-spectrometry proteomics.

This work was supported by the following grants (ZonMW-TOP BECAUSE (91216064), to M.V and L.N.C. ZonMW-PSIDER Brainmodel (10250022110003) to M.V and L.N.C., NWO-Dutch Research Agenda (NWA) NewTDec (1160.18.200) to M.V and L.N.C.), NWO Gravitation program grant BRAINSCAPES (NWO 024.004.012) to M.V. and ZonMW-OPEN (9120232310098) from the Netherlands Organisation for Scientific Research (NWO) to M.V. and L.N.C.

## Author contributions

M.E.W., and J.M.C.M. designed and produced the PiggyBac construct

T.W.v.V, M.A.v.B., K.I.M., R.F.T., A.B.S., R.E.v.K, M.V. and L.N.C. designed the experiments T.W.v.V, M.A.v.B. and J.S. performed and maintained neuronal inductions and cultures

T.W.v.V M.A.v.B. performed the patch-clamp experiments with contributions of J.S. to Fig. 1 and C.F. to Fig. S2

T.W.v.V, M.A.v.B.,. performed the immunocytochemistry experiments with contributions of J.S to Fig. 1

K.I.M., and F.K. performed the proteomics experiments and analysis

T.W.V.V, M.A.v.B., K.I.M., R.E.v.K, M.V. and L.N.C. designed figures and wrote the manuscript with input from all authors.

## Supplementary information

Supplementary information will be made available with the manuscript.

## Extended Methods

### Key resources table

| REAGENT or RESOURCE | SOURCE | IDENTIFIER |
| --- | --- | --- |
| Antibodies |  |  |
| MAP2 | Merck Millipore | (Millipore Cat# AB5543, RRID:AB_571049) |
| VGAT | Synaptic Systems | (Synaptic Systems Cat# 131 002, RRID:AB_887871) |
| VGLUT1 | Antibodies Inc. | (Antibodies Incorporated Cat# 75-066, RRID:AB_2187693) |
| VGLUT2 | Antibodies Inc. | (Antibodies Incorporated Cat# 75-067, RRID:AB_2239153) |
| NeuN | Merck Millipore | (Millipore Cat# ABN78, RRID:AB_10807945) |
| Parvalbumin | Synaptic Systems | (Synaptic Systems Cat# 195 004, RRID:AB_2156476) |
| **Bacterial and virus strains** |  |  |
| FUW-TetO-*Ascl1*-T2A-puro | Addgene | RRID:Addgene_97329 |
| FUW-TetO-*Dlx2* -IRES-hygro | Addgene | RRID:Addgene_97330 |
| FUW-M2rtTA | Addgene | RRID:Addgene_20342 |
| **Deposited data** |  |  |
| PRIDE dataset " One-step induction of human GABAergic neurons promotes presynaptic development & synapse maturation" | This study | PXD064163 |
| Transcriptomic cytoarchitecture reveals principles of human neocortex organization | Jorstad, et al.<sup>40</sup> | RRID:SCR_016152 |
| **Experimental models: Cell lines** |  |  |
| BiONI010-C-10 | EBiSC | RRID:CVCL_RM84 |
| BioNi010-C-13 | EBiSC | RRID:CVCL_RF90 |
| Kolf2.1J | Jackson Laboratory | RRID:CVCL_B5P3 |
| GM23973 | Coriell | RRID:CVCL_BW42 |
| **Recombinant DNA** |  |  |
| PB-TO-ASCL1-DLX2 | This study | RRID:Addgene_182307 |
| Transposase | This study | N/A |
| **Software and algorithms** |  |  |
| DIA-NN 1.8.1 | Demichev, et al<br>(2020,2022)<sup>154,155</sup> |  |
| MS-DAP 1.0.5 | Koopmans, et al.<br>2023<sup>150</sup> |  |
| DEqMS | Zhu, et al, 2020<sup>151</sup> |  |
| GOAT | Miedema, et al.<br>2022<sup>152</sup> |  |
| SynGO | Koopmans, et al.<br>2019<sup>153</sup> |  |

### iPSCs origin and culture

Unless stated otherwise, the BiONi010-C-10 stem cell line (EBISC, RRID:CVCL_RM84) was used for all experiments in this study. For glutamatergic induction, a gene-edited variant of the same line containing an integrated NGN2-casette was used (EBISC, RRID:CVCL_RF90). As a genetically independent control the KOLF2.1J (JAX, JiPSC1000, RRID:CVCL_B5P3) and GM23973 (Coriell, RRID:CVCL_BW42) lines were used. All iPSCs were maintained on Matrigel (Cornig) or Geltrex (Fisher Scientific)-coated dishes in Essential 8 (Gibco) or Essential 8 Flex (Gibco) medium and passaged with EDTA and replated in the culture medium supplemented with ROCK inhibitor Y27632 (5 µM, Tetu bio).

### Lentiviral infection

Ultrahigh titre lentiviral particles were generated by Alstem (Richmond, CA, USA) from FUW-M2rtTA (Addgene, #20342, RRID:Addgene_20342), FUW-TetO-*Ascl1*-T2A-puro (Addgene #97329, RRID:Addgene_97329), and FUW-TetO-*Dlx2* −IRES-hygro (Addgene #97330, RRID:Addgene_97330). For *Ascl1*+Forskolin, iPSCs were infected with only FUW-M2rtTA and TetO-*Ascl1*-Puro, for *Ascl1 + Dlx2* induction, iPSCs were infected with all three lentiviruses. For both lentiviral induction methods, 24 hours after infection, puromycin (1 µg/ml) selection was performed. For lentiviral *Ascl1* + *Dlx2* induction, additional hygromycin (75 µg/ml) selection was performed. For both selection regimes an uninfected control well was included to confirm successful selection before proceeding towards induction.

### PiggyBac transfection

A more detailed protocol is found at the end of this document, see <u>protocol 1</u>. To prepare for PiggyBac transfection, iPSCs were released with accutase and plated as single cells on a Matrigel-coated 6-well plate in E8 + ROCK Inhibitor Y27632 at a density of 1500k per well. Cells were placed in the incubator to adhere for 1 to 2 hours. A total of 3 µg of DNA containing the *Ascl1 + Dlx2* PiggyBac transposon (Presented here; available on Addgene, 182307, RRID:Addgene_182307) and transposase constructs (extended data 1) at a 1:3 molar ratio was prepared in 100 µL of opti-MEM (GIBCO, 31985062). To another 100 µL op opti-MEM 5 µL of Lipofectamine STEM (Invitrogen, STEM00001) was added. Opti-MEM preparations were quickly mixed by pipetting, after which the mixture was incubated at RT for 15 to 30 minutes during which cells received fresh E8 + ROCK Inhibitor Y27632 to wash away non-adherent cells. The opti-MEM solution was then added to the cells in a dropwise fashion. 24 hours post transfection, selection with puromycin (1 µg/ml) was initiated to select only for cells containing the transposon, and selection was repeated every 3-7 days.

### PB-AD GABAergic neuronal induction

A more detailed protocol is found at the end of this document, see <u>protocol 2</u>. PB-AD positive iPSCs were released with Accutase (Merck) on day post induction (DPI) 0 and replated with at an approximate density of 3.500 cells/cm^2^ in N2 medium (DMEM/F12 medium, Life technologies) containing Glutamax (200 mM, Life Technologies), dextrose (0.3%, Life Technologies) N2 supplement B (StemCell Technologies), Pen/Strep (0.1%, Gibco), ROCK inhibitor Y27632 (10 µM, Tetubio) and doxycycline hyclate (2 µg/mL) onto Matrigel (Cornig) or Geltrex (Fisher Scientific) coated dishes. On DPI 1, medium was refreshed using the same composition without ROCK inhibitor Y27632. On DPI 2, the medium was refreshed once more with the same composition as on DPI 1, with the addition of 5-Fluoro-2’-deoxyuridine (FUDR) (10 µM, Sigma-Aldrich) or Cytosine β-D-arabinofuranoside (AraC, 20 µM, Sigma-Aldrich) to remove proliferative cells. On DPI 3, neurons were released with accutase, pelleted, and resuspended and plated in Neurobasal medium (Life Technologies) supplemented with Glutamax (200 mM), B27 (0.1%) Pen-Strep (0.1%), fetal bovine serum (0.5%), BDNF (10 ng/ml, Peprotech), CNTF (10ng/ml, Peprotech), GDNF (10 ng/ml, Peprotech) and doxycycline hyclate (2 µg/mL; Sigma-Aldrich). Neurons were plated to the required respective preparations (see *Autaptic culture* and *Network culture*).

### Glutamatergic neuronal induction

For glutamatergic induction, Bioni010-C-13 iPSCs were released with Accutase (Merck) on DPI 0 and replated with at an approximate density of 3.500 cells/cm^2^ in N2 medium (DMEM/F12 medium, Life technologies) containing Glutamax (200 mM, Life Technologies), dextrose (0.3%, Life Technologies) N2 supplement B (StemCell Technologies), Pen/Strep (0.1%, Gibco) and doxycycline hyclate (2 µg/mL; Sigma-Aldrich) combined with dual-SMAD inhibitors LDN193189 (100 nM, Stemgent), SB431542 (10 µM, Tocris Bioscience), and XAV939 (2 µM, Stemgent) NT3 (10 ng/ml, Peprotech) and doxycycline hyclate (2 µg/mL) onto Matrigel (Cornig) or Geltrex (Fisher Scientific) coated dishes. On DPI 1, medium was refreshed using the same composition without ROCK inhibitor Y27632. On DPI 2, the medium was refreshed once more with the same composition as on day 2, with the addition of 5-Fluoro-2’-deoxyuridine (FUDR) (10 µM, Sigma-Aldrich) or Cytosine β-D-arabinofuranoside (AraC, 20 µM, Sigma-Aldrich) to remove proliferative cells. On DPI 3, neurons were released with accutase, pelleted, resuspended and plated in Neurobasal medium (Life Technologies) supplemented with Glutamax (200 mM), B27 (0.1%) Pen-Strep (0.1%), foetal bovine serum (0.5%), BDNF (10 ng/ml, Peprotech), CNTF (10ng/ml, Peprotech), GDNF (10 ng/ml, Peprotech) and doxycycline hyclate (2 µg/mL; Sigma-Aldrich). Neurons were plated to the required respective preparations (see *Autaptic culture* and *Network culture*).

### Lentiviral GABAergic induction

In general, lentiviral GABAergic induction protocols followed a similar workflow as PB-AD induction. To better adhere to established protocols.^14,15^, the medium composition differed at certain steps (Figure 1H). For Ascl1 + Forskolin induction, from DPI 0 onwards, Forskolin (10 µM, Merck) was added to the medium up to DPI 14. Furthermore, from DPI0 onwards non-essential amino acids (1:100, Sigma) were added up to day DPI 14. BDNF (10 ng/ml) and NT3 (10 ng/ml) were present from DPI1 onwards. For lentiviral *Ascl1* + *Dlx2* induction BDNF (10 ng/ml) and NT3 (10 ng/ml) were added from DPI 0 onward, as well. Furthermore, laminin (1µg/ml, Sigma-Aldrich), cyclic AMP (1 µM, Sigma-Aldrich), and Ascorbic Acid (200 µM, Sigma-Aldrich), were added to the medium at all subsequent refreshments to improve viability.

### Autaptic culture

Neurons were plated at a density of ∼ 400 cells / cm^2^ on micro-islands prepared from rat glial cells plated on agarose-coated coverslips stamped with a custom stamp in Poly-D-Lysine (0.1 mg/ml, Sigma), rat tail collagen (0.7 mg/ml, Corning) and Fibronectin (100 µg/ml, Corning) in acetic acid (10 mM, sigma). Glial islands were prepared approximately 7 days before plating neurons and were treated with AraC (20 µM, Sigma-Aldrich) before plating neurons. Autaptic cultures were refreshed every 7 days with a ∼50% media refreshment, with the same medium as used for plating neurons, but omitting doxycycline hyclate (except for Figure 1E-G, where doxycycline hyclate was added up to DPI 14).

### Network culture

Neurons were plated at a density of ∼ 7000 cells / cm^2^ on coverslips covered with a glial-monolayer prepared from rat astrocytes a week in advance and treated with Cytosine β-D-arabinofuranoside (20 µM, Sigma-Aldrich) 24-48 hours before plating neurons. Networks cultures were refreshed with a ∼50% media refreshment every 3-4 days, with the same medium as used for plating neurons, but omitting doxycycline hyclate.

### Electrophysiology

Whole-cell voltage-clamp recordings (V^m^ = −70 mV) were performed on autaptic neurons, with borosilicate glass pipettes (2–5MΩ) filled with (in mM) 136 KCl, 17.8 HEPES, 1 EGTA, 0.6 MgCl^2^, 4 ATP-Mg, 0.3 GTP-Na, 12 phosphocreatine-K^2^ and 50 units/mL phosphocreatine kinase (pH = 7.30, 290 mOsmol). The external solution contained in mM: 10 HEPES, 10 glucose, 140 NaCl, 2.4 KCl, 4 MgCl^2^, and 4 CaCl^2^ (pH = 7.30, 290 mOsmol). Experiments were performed at room temperature (∼22℃ using a MultiClamp700B amplifier (Axon Instruments) and Digidata 1440A digitizer under control of Clampex 10 software (Molecular devices). Only cells with an access resistance < 15MΩ (80% compensated) and leak current of <300 pA were included. Synaptic responses were elicited by 0.5 ms depolarization to 30 mV.

For GABA puffs, perforated patch recordings were produced by filling the patch pipette (in mM) 140 KCl, 10 HEPES, gramicidin (100 µg/ml) and allowing cells to perforate for 15-30 minutes after reaching a GΩ seal to < 100 MΩ series resistance. Puff pipettes filled with 100 µM GABA in ACSF were positioned near the soma, at approximately 25 µm distance. 100 ms puffs were applied at −110,-90,-70,-50,-30 and −10 mV holding voltages to induce GABAergic currents. Cells were discarded if a sudden swift decrease in resistance was observed during perforation, or if the observed reversal potential of Cl^-^ was ± 0 mV, which indicates exchange of the patch-pipette solution with the intracellular solution.

### Electrophysiology; pharmacology

To Block GABA-currents, picrotoxin (100 µM, Hello Bio) was used. A 100 mM stock was prepared in DMSO and was added to the bath to reach the final concentration. To block glutamatergic currents, AP-V (100 µM, R&D systems) and DNQX (40 µM, Tocris) were used. For DNQX, A 1000x of stock of each was prepared in DMSO. For AP-V, the stock was prepared in water.

### Electrophysiology; analysis electrophysiological data

Offline analysis was performed using custom-written programs^156^ in Matlab (Mathworks). Experimental conditions were blinded during analysis. In all figures, stimulation artefacts have been removed. For evoked release during 20 Hz stimulation, back-extrapolation of linear fits from the last 20 pulses of the cumulative charge was used to estimate the RRP (y-intercept).^157^

### Immunocytochemistry and morphological analysis

Neurons were fixed in ice cold 100% methanol (VWR) for 15 minutes at 4°C, blocked with 2.5% normal goat serum (NGS, Life Technologies) and 0.1% Triton X-100 in PBS for 45 minutes and incubated with primary antibodies in PBS with 2.5% NGS and 0.1% Triton X-100 overnight at 4°C. The following primary antibodies were used: chicken anti-MAP2 (1:1000, Merck AB5543, RRID:AB_571049); rabbit anti-VGAT (1:2000, SySy #131 002, RRID:AB_887871), mouse anti-VGlut1 (1:250, NeuroMab clone N28/9, Antibodies inc. 75-066, RRID:AB_2187693), mouse anti-VGlut2 (1:250, NeuroMab clone N29/29, Antibodies inc. 75-067, RRID:AB_2239153). For Rabbit anti-NeuN (1:500, Merck ABN78, RRID:AB_10807945) and anti-Parvalbumin (1:500, SySy #195 004, RRID:AB_2156476) staining neurons were fixed with 3.7% PFA for 15 minutes at room temperature, instead of methanol fixation. After washing with PBS the next day, cells were incubated for 1 h at room temperature with secondary antibodies conjugated to Alexa dyes (1:1000, Invitrogen Molecular Probes), washed again and mounted with DABCO-Mowiol (Invitrogen). Images were acquired at 1024×1024 pixels on a Nikon Eclipse Ti confocal laser scanning microscope (40× objective; NA 1.3) with NIS-Elements software. Z-stacks were produced at a 0.5 µm step-size until no more puncta and/or dendrites were in focus. Confocal settings were kept constant for all scans within an experimental week. Analysis of neuronal morphology was performed on maximum-intensity projections using the automated image analysis routine SynD.^147^ for autaptic neurons and a custom ImageJ workflows for network analysis, in either case, for VGAT and VGLUT puncta counts, only puncta on dendrites were counted.

### Statistics of electrophysiological and morphological data

Statistical testing was performed in MATLAB and GraphPad Prism For descriptive statistics (Figure 1) the likelihood of normality vs lognormality was calculated, which revealed that the data was lognormally disturbed. Therefore the geometric mean and geometric SD factor were calculated and reported. In autaptic development timelines (Figure 4&5), no data was normally distributed. Therefore, to test general developmental trajectories, nonparametric Spearman correlations were computed with a two-tailed test for significance and a 95% confidence interval. For timepoint comparisons, Kruskal-Wallis ANOVAs were performed with Dunn’s multiple comparisons test. Pairwise comparisons were made between 4 and 6, 6 and 8, and 8 and 10 WPI. #, ##, ###, #### and *ns* represent results of spearman correlation results, *, **, ***, **** and ns represent Kruskal-Wallis ANOVA results. *ns* and ns represent non-significant results. Single, double, triple and quadruple marks represent (corrected) p <0.05, p <0.01, p <0.001 and p <0.0001, respectively. GABA-puff responses (figure 6O) were fitted using a first-order polynomial linear fit and an extra sum-of-squares F-test was used to determine whether GABA responses were identical throughout development. In plots, unless stated otherwise, the lines in boxplots represent the median value. Crosses represent the mean value. Boxes represent 50% off the values. Circular markers represent individual observations. Quantifications are available in the supplementary data, and each figure contains n (number of cells) and N (number of inductions) values per parameter in either the relevant graph or typical example.

### Mass spectrometry; Cell pellet harvesting for MS

Cells were washed three times with 1 ml sterile Dulbecco’s PBS (Gibco), whereof first twice at 37°C and finally once at 4 °C. Cultures were kept on ice, scraped off using VWR cell lifters, and harvested in 1.5 ml protein LoBind Eppendorf tubes (VWR). Pellets were obtained by centrifugation at 1,000 × g for 5 minutes at 4 °C. Supernatant was removed and pellets were kept at −80 °C before subsequent sample preparation.

### Mass spectrometry; MS sample preparation

MS sample preparation was performed following the DNA micro spin column suspension trapping protocol^148^, with some minor adaptations. Cell pellets were extracted and reduced in 6% SDS buffer, containing 50 mM Tris-HCl (pH 8) and 5 mM tris (2-carboxyethyl)phosphine (TCEP), by incubation at 55°C for 15 minutes in a thermomixer set to 1400 RPM. Free sulfhydryl groups were alkylated by incubation with 20 mM methyl methanethiosulfonate (MMTS; 200 mM stock solution) for 15 minutes at RT. Insoluble debris was cleared by centrifugation at 10000g for 1 minute at RT and the supernatant was transferred to a new tube (Eppendorf). Next, samples were acidified by addition of 1.1% phosphoric acid (12% stock solution), mixed with six volumes of binding/washing buffer (90% methanol and 100mM Tris-HCl, pH 8) and loaded onto a plasmid DNA micro column (HiPure from Magen Biotechnology). The protein particulate was retained on the column upon centrifugation at 1400x g for 1 minute and the columns were washed four times with binding/washing buffer. Columns were transferred to a new tube (Eppendorf), supplemented with 500 ng Trypsin/Lys-C (Promega) in 50 mM NH4HCO3 and incubated overnight at 37°C in a humidified incubator. Tryptic peptides were eluted and pooled by subsequent addition of 50 mM NH4HCO3, 0.1% formic acid and 0.1% formic acid in acetonitrile. Collected peptides were dried by SpeedVac and stored at −80°C.

### Mass spectrometry; LC-MS Analysis

Each sample of tryptic digest was redissolved in 0.1% formic acid and the peptide concentration was determined by tryptophan-fluorescence assay^158^; 75 ng of peptide was loaded onto an Evotip Pure (Evosep). Peptide samples were separated by standardized 30 samples per day method on the Evosep One liquid chromatography system, using a 15 cm × 150 µm reverse-phase column packed with 1.5 µm C18-beads (EV1137 from Evosep) connected to a 20 µm ID ZDV emitter (Bruker Daltonics). Peptides were electro-sprayed into the timsTOF Pro 2 mass spectrometer (Bruker Daltonics) equipped with CaptiveSpray source and measured with the following settings: Scan range 100-1700 m/z, ion mobility 0.65 to 1.5 Vs/cm2, ramp time 100 ms, accumulation time 100 ms, and collision energy decreasing linearly with inverse ion mobility from 59 eV at 1.6 Vs/cm2 to 20 eV at 0.6 Vs/cm2. Operating in dia-PASEF mode, each cycle took 1.38 s and consisted of 1 MS1 full scan and 12 dia-PASEF scans. Each dia-PASEF scan contained two isolation windows, in total covering 300-1200 m/z and ion mobility 0.65 to 1.50 Vs/cm2. Dia-PASEF window placement was optimized using the py-diAID tool^159^ Ion mobility was auto calibrated at the start of each sample (calibrant m/z, 1/K0: 622.029, 0.992 Vs/cm2; 922.010, 1.199 Vs/cm2; 1221.991, 1.393 Vs/cm2).

### Mass spectrometry; MS data Analysis

DIA-PASEF raw data were processed with DIA-NN 1.8.1.^154,155^ An in-silico spectral library was generated from the uniprot human reference proteome (SwissProt and TrEMBL, canonical and additional isoforms, release 2023-2) using Trypsin/P digestion and at most 1 missed cleavage. Fixed modification was set to beta-methylthiolation (C) and variable modifications were oxidation (M) and N-term M excision (at most 1 per peptide). Peptide length was set to 7-30, precursor charge range was set to 2-4, precursor m/z was limited to 280–1220, both MS1 and MS2 mass accuracy were set to 15 ppm, scan window was set to 9, double-pass-mode and match-between-runs were enabled. Protein identifiers (isoforms) were used for protein inference. All other settings were left as default.

MS-DAP 1.0.5.^150^ was used for downstream differential abundance analyses using the DIA-NN results. Filtering and normalization was applied to respective samples per statistical contrast. Peptide-level filtering was configured to retain only peptides identified in at least 4 samples in both experimental conditions (within current contrast) and only proteins with 2 valid peptides were retained. Next, peptide abundance values were normalized by minimizing the mode log^2^ (FC) between sample pairs (MWMB algorithm), followed by protein-level mode-between normalization. The DEqMS^151^ algorithm was used for statistical analysis and resulting p-values were adjusted for multiple testing using the Benjamini–Hochberg False Discovery Rate (FDR) procedure. To investigate proteins with strong differences in detection rates between conditions, e.g. consistently detected in one condition but (near) absent in another, the MS-DAP differential detection test was applied to all proteins that were observed with at least 2 peptides in at least 4 samples in either condition (i.e. confidently detected proteins). Using the default MS-DAP workflow, we merged differential detection (DD) results into the DEqMS result table for the subset of proteins that had both an absolute DD z-score of at least 6 and a smaller adjusted p-value in DD (or no p-value from DEqMS due to a lack of data points in either condition).

### Mass spectrometry; Expression-weighted cell type enrichment analysis

Cell type enrichment analysis gives insight into the cell types that potentially drive differential expression. EWCE uses single-cell RNA-sequencing data to generate a probability distribution associated with a gene list having an average level of expression within a cell^41^ Although it’s based on RNA expression levels, it can also be applied to proteomics data.^160^ We used published single nucleus RNA-sequencing data (49,495 nuclei) from human cortical brain regions – including middle temporal gyrus, anterior cingulate cortex, primary visual cortex, primary motor cortex, primary somatosensory cortex and primary auditory cortex – published by Jorstad et al.^40^ For cell type annotations we used the cell type annotations: class (e.g., GABAergic neurons) and subclass (e.g. SST^+^ interneurons) for first and second level EWCE analysis, respectively. Bootstrap enrichment tests were performed using 10,000 reps to look whether gene sets (i.e., foreground) were significantly enriched for a cell class or subclass relative to the background (i.e., all detected proteins included in that contrast). Since we focused on neuronally-enriched proteins or SYNGO-annotated synaptic proteins, we excluded non-neuronal enrichment scores.

EWCE calculates cell type-specificity scores (CTSS) for all genes, which is the percentage of mean expression per cell type relative to the sum of the mean expressions of all cell types. We used the CTSS to predict for each protein the cell type the protein probably is derived from. We only included CTSS from genes that passed mRNA detection threshold (mean expression >4 in at least one cell class). Cell type predictions included neuronal (neuronal CTSS > 70%), GABAergic neurons (GABA CTSS > 0.8 and glutamatergic CTSS < 0.1), glutamatergic neurons (glutamatergic CTSS > 0.8 and GABAergic CTSS < 0.1). Cell type predictions were labelled in volcano plots. We also used the EWCE-calculated mean mRNA expression for each cell type to estimate the differential expression of genes in GABAergic vs glutamatergic neurons, which was used in the SYNGO manual colour plots (Figure 6H)).

### Mass spectrometry; Heatmaps and line plots

We used the mean log^2^-transformed normalized protein intensities per group (i.e., PBAD 4 WPI) and plotted these values using line plots in RStudio.^161^ using ggplot^162^ The relative normalized protein intensities (log^2^ (FC)) was used in heatmaps, which were produced using Morpheus. To map these proteins to predicted cell types we also show EWCE-calculated cell type specificity scores. All heatmaps visualizations are with fixed colour scaling (i.e., not relative colour scaling per gene), as indicated by heatmap legend.

### Mass spectrometry; Gene-set enrichment analysis using GOAT and over-representation analysis using SYNGO

We used the output of the differential abundance analysis using MSDAP (PB-AD 10 vs 4 WPI and NGN2 10 vs 4 WPI) as input for gene-set enrichment analysis using GOAT. The GOAT online v1.0.2.^152^ was used to perform gene set enrichment analyses using gene sets from the SynGO knowledgebase v1.2^153^ Effect size-derived gene scores (i.e., log^2^ (FC)) were used to test for enriched gene sets that contained at least 10 and at most 1500 genes (or 50% of the gene list, whichever was smaller) that overlapped with the input gene list. Multiple testing correction was independently applied per gene set “source” (i.e. SYNGO_CC, SYNGO_BP) using Bonferroni adjustment and all p-values were again adjusted using Bonferroni correction to account for 2 separate tests across “sources”. The significance threshold for adjusted p-values was set to 0.05. SYNGO over-representation analysis was performed using the same MSDAP output. Foreground was defined as upregulated proteins per contrast (p-adjusted < 0.05 and log^2^ (FC) > 0.1), and the background contained all tested genes for that contrast. SynGO release is dataset version: 20231201, and default settings were used. Ontologies are significantly enriched at 1% FDR, while including terms containing at least three matching input genes.

### Protocol 1: PiggyBac transfection

#### Introduction

In this protocol it is described how IPSCs are transfected with the transposon and transposase to allow for neuronal induction of PB-AD neurons.

#### Materials

- Geltrex (Cornig) or Matrigel (Fisher Scientific)
- Essential 8 (Gibco) or Essential 8 Flex (Gibco)
- Accutase (Millipore; SCR005)
- DMEM-F12 (Life technologies)
- opti-MEM (GIBCO, 31985062)
- Lipofectamine STEM (Invitrogen, STEM00001)
- Rock Inhibitor Y27632 (Tetu-Bio, T1725) or CEPT cocktail (chroman 1 (Mechem Expresss, HY-15392), emricasan (Selleckchem, S7775), polyamine (Sigma, P8483), and trans-ISRIB (Tocris, 5284))
- Doxycycline hyclate 2 µg/mL (Sigma, D9891)
- Puromycin (Sanbio, 13884-25)
- Piggybac transposase (see supplementary plasmid map)

- Distribution of the transposase through Addgene is not possible. We advise synthesis inhouse or commercially using the supplied plasmid map.
- *Ascl1 + Dlx2* PiggyBac transposon (Addgene, 182307, RRID:Addgene_182307)

#### Protocol

- Coat your plates with Geltrex or Matrigel according to manufacturer instructions. Culture IPSCs in 1×6w until 70-90% confluent.
- Prepare a fresh coated 1×6w for plating infected cells. Prewarm Essential 8 or Essential 8 Flex + Rock Inhibitor Y27632 (RI, 10 µm) or CEPT (50 nM, 5 µM, 1×, 07. µM respectively).

- Some IPSC lines are more easily transfected then others. If severe cell-death or bad transfection efficiency are observed, CEPT cocktail can improve this.
- Aspirate the medium of the IPSCs and wash 1× with PBS.
- Aspirate medium and add 350 µl accutase to 1×6w IPSCs. Incubate at 37 °C For approx. 5-10 minutes or until cells detach.
- Meanwhile, aspirate the coating medium from a new plate and add Essential 8 or Essential 8 Flex + RI or CEPT.
- Once cells have detached, collect cells with 4.5 ml DMEM + RI and spin down 5 minutes at 1000 RPM / 180 RCF.
- Aspirate the supernatant and resuspend cells in 1 ml of Essential 8 or Essential 8 Flex + RI or CEPT cocktail.
- Count cells and plate 1000 to 1500k cells per 1×6w well, 500 to 750k cells per 1×12w or 200 to 300k per 1×24w.
- Let cells adhere for 1 to 2 hours at 37 °C/5% CO^2.^
- Per transfection, prepare the mixes as stated in table 1. Give the Opti-MEM a few minutes to warm to RT before adding the other components to each mix.

Table: Mixing ratio for transfection of piggyBac constructs

| 6 well |  |  | 12 well |  |  | 24 well |  |  |
| --- | --- | --- | --- | --- | --- | --- | --- | --- |
|  | Mix 1 | Mix 2 |  | Mix 1 | Mix 2 |  | Mix 1 | Mix 2 |
| Opti-MEM | 100 µl | 100 µl | Opti-MEM | 50 µl | 50 µl | Opti-MEM | 20 µl | 20 µl |
| DNA | 3 µg |  | DNA | 1.5 µg |  | DNA | 0.6 µl |  |
| Lipofectamine stem |  | 5 µl | Lipofectamine stem |  | 2.5 µl | Lipofectamine stem |  | 1 µl |

- For DNA, mix the transposase and transposon at a molar ratio of 1:3.
- Combine mix 1 and 2 and mix well by pipetting up and down. Incubate transfection mix for 15 to 30 minutes at RT.
- Replace Essential 8 or Essential 8 Flex + RI or CEPT on the cells during this incubation step, to wash away any non-adherent cells and debris.
- Add the mixture to the cells in a dropwise fashion, evenly distributed throughout the well. Immediately swirl the plate to ensure even distribution, and return plate to 37 °C/5% CO^2^.
- Over the following 24 to 72 hours, BFP expression should appear. Culture cells to 70-90% confluency or for 1 week before splitting if confluency isn’t reached.
- Once cells are 50-70% confluent, add puromycin for selection. This selection regime can be started 48h after transfection, if cells divide quickly, or later. The required concentration can vary between cell-lines. For most lines, 1 µg/ml puromycin for a minimum of two days is recommended.

- Take along a well of untransfected IPSCs of the same genotype as a control and only stop selection once no cells survive in this well.
- Expand selected cells and use as needed. Repeated antibiotic selection should be performed weekly to ensure stable integration of the transposon is maintained.

### Protocol 2: PB-AD GABAergic neuronal induction

#### Materials

- Matrigel (Cornig), Geltrex (Fisher Scientific)
- Accutase (Millipore; SCR005)
- Rock Inhibitor Y27632 10 µM (Tetu-Bio, T1725) (RI)
- Doxycycline hyclate 2 µg/mL (Sigma, D9891) (Dox)
- FUDR (Sigma, F0503) 10 µM **or** AraC (Sigma, C1768) 20 µM

- AraC more efficiently selects only postmitotic cells and is recommended, but if lentiviral infection after induction is needed FUDR selection allows for higher infection efficiency after selection.
- Neurotrophic Factors

- BDNF (Stemcell Technologies; 78005.3) - 10 ng/ml
- CNTF (Peprotech; 450-13) - 10 ng/ml
- GDNF (Stemcell Technologies; 78058.3) - 10 ng/ml
- DMEM (VWR; 392-0415P)

- foetal bovine serum (2%, Life Tech; 10270106)
- N2 Medium

- DMEM/F12 medium (Life Technologies)
- Glutamax (200 mM, Life Technologies)
- dextrose (0.3%, Life Technologies)
- N2 supplement B (StemCell Technologies)
- Pen/Strep (0.1%, Gibco)
- Neurobasal Medium

- Neurobasal medium (Life Technologies)
- Dextrose (Life Tech; A2494001) – 0.3% final concentration – use 1:67
- Glutamax (200 mM)
- NEAA (Life Tech; 11350912) - 1:200
- B27 (0.1%)
- Pen-Strep (0.1%)
- foetal bovine serum (Life Tech; 10270106, 0.5%)

### Protocol

**Day 0: Start induction**

1 Per 1×6w IPSCs coat 1× 10cm dish with Geltrex or Matrigel according to manufacturer instructions. Prewarm 10 ml of N2 + RI + Dox, 5 ml of DMEM + 2% FBS per 1×6w to induce, and 5 ml PBS.
2 Remove media from IPSCs and wash 1× with warm PBS.
3 Add 300 µl accutase per 6-well.
4 Incubate for 5 minutes at 37°C.
5 Add 3 ml DMEM and pipet up and down to collect cells.
6 Pellet for 5 minutes at 1000 RPM / 180 RCF.
7 Resuspend pellet in 1 ml N2 medium with RI + Dox.
8 Plate cells as single-cells in a 10 cm dish with 9 ml N2 medium with RI + Dox.

**Day 1: Repeat Doxycycline and remove RI**

9 The following day, replace 100% of the medium with N2 + Dox

**Day 2: Repeat Doxycycline and select postmitotic cells**

10 The following day, replace 100% of the medium with N2 + Dox + FUDR **or** N2 + Dox + AraC

**Day 3: Replate neurons to target vessel**

11 Prepare Neurobasal medium + DOX + CNTF + GDNF + BDNF (if using FUDR, + FUDR)
12 When media is warm, transfer it to prepared plates. Induced GABAergic neurons thrive in co-culture with glia. Neurons can be replated to coverslips pre-seeded with glia, or be mixed with glia and plated simultaneously.
13 Wash induced neurons 1× with PBS.
14 Add accutase to your induced neurons (1.5 ml in 10 cm dish).
15 Incubate for 5 minutes at 37°C or until cells are released.
16 Add 3 ml of DMEM + 2% FBS and pipette up and down to collect cells.
17 For 1×10 cm dish, use 13 ml of medium for collection.
18 Pellet cells for 5 min at 1000 RPM / 180 RCF.
19 Resuspend the pellet in 1 ml NBM + Dox + BDNF + CNTF + GDNF and count cells.
20 Plate neurons at your required density.

#### Feeding

21 Feeding rates depend on experimental conditions. For the experiments as described here, low density (autaptic) cultures are fed once a week. Higher density cultures are fed twice a week. In all cases, a ∼50% medium refreshment is performed. Media composition is the same as the glia-transfer medium, but AraC/FUDR is omitted.

## Material availability

PiggyBac *Ascl1*-*Dlx2* construct is available from Addgene <u>#182307</u>. The mass spectrometry proteomics data have been deposited to the ProteomeXchange Consortium via the PRIDE partner repository.^163^ with the dataset identifier PXD064163.

## Supplementary figures and table

**Figure S1.**
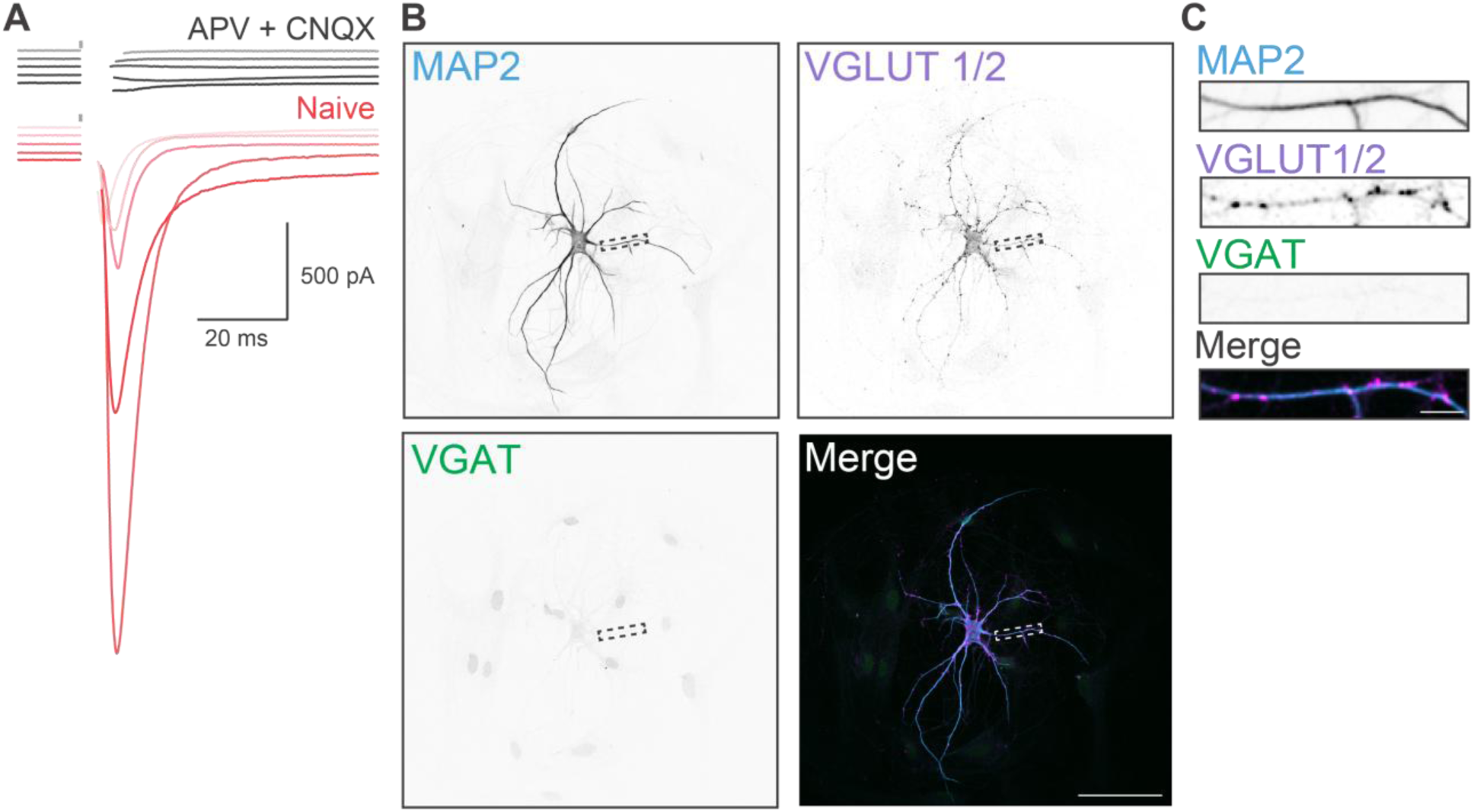
NGN2 neurons produce glutamatergic transmission. **A** Voltage clamp recordings of 6 WPI NGN2 autaptic neurons after stimulation, in normal ACSF (Naive) and in presence of glutamatergic receptor blockers (APV + CNQX). Synaptic transmission was fully abolished by blockers, illustrating that NGN2 neurons are glutamatergic. **B-C** Examples of immunocytochemistry for MAP2, VGAT and VGLUT1+2 performed on NGN2 autaptic neurons, with zoom in on a length of dendrite (**C**). Scale bar = 100 µm, zoom in scale bar = 10 µm

**Figure S2.**
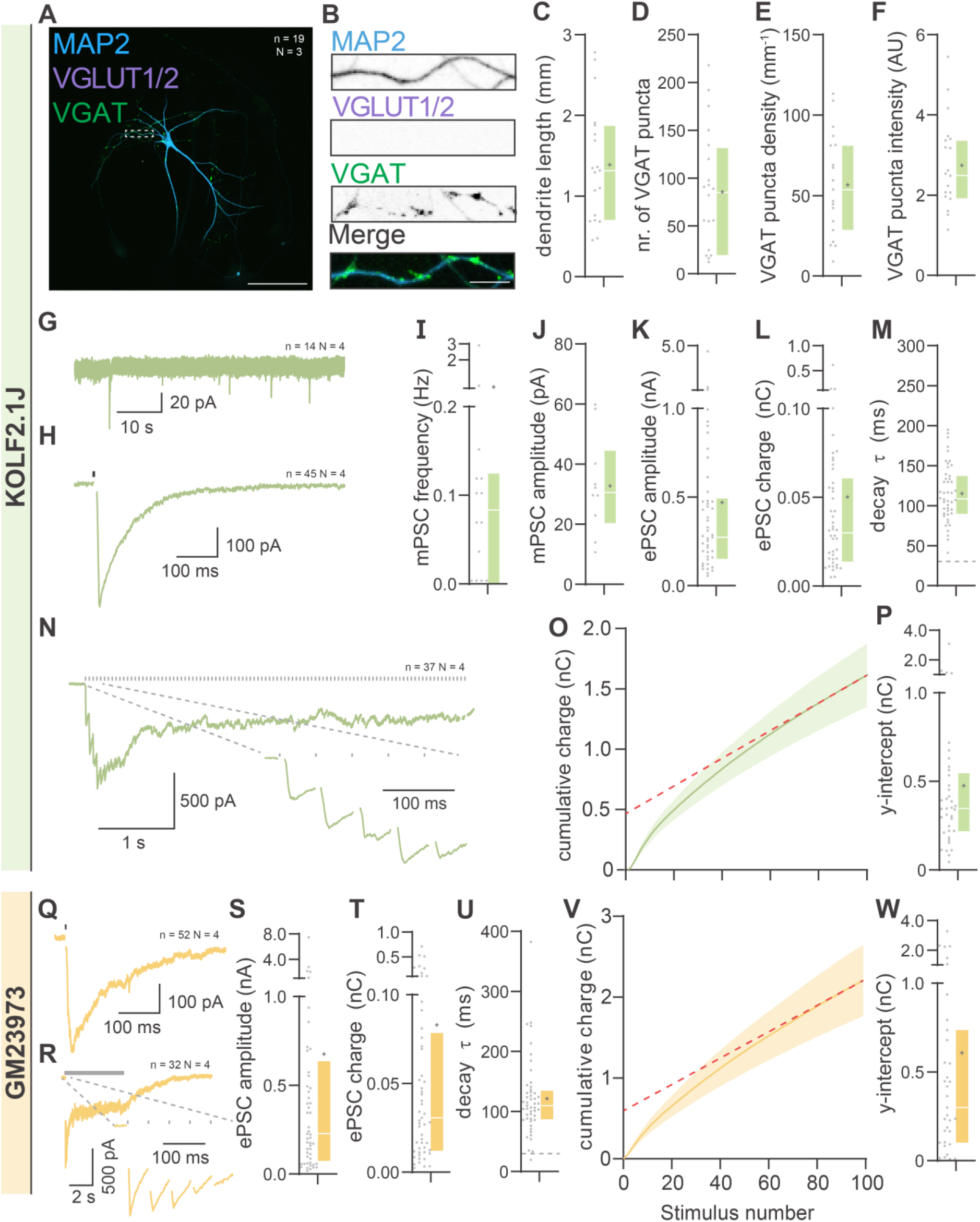
PiggyBac Ascl1/Dlx2 induction reliably generates GABAergic neurons in the KOLF2.1J and GM23973 line. **A** Representative images of PiggyBac (PB) Ascl1/Dlx2 (AD) neurons induced from KOLF1.2J iPSCs at 7 weeks post induction (WPI), stained for MAP2 as dendritic marker, and VGAT and VGLUT1/2 as GABAergic and glutamatergic markers, respectively. Scale bar: 100 µm **B** Zoom-in on dendrite to illustrate punctate VGAT staining. Scale bar: 10 µm **C.** Total dendrite length and (**D**) number of GABAergic synapses per neuron. **E** Synaptic density and intensity (**F**). **G** Representative mPSC trace and (**H**) representative ePSC trace from PB-AD neurons induced from the KOLF2.1J line. **I** Frequency and (**J**) amplitude of miPSCs. **K** Amplitude, (**L**) charge and (**M**) decay time constant of iPSCs in PB-AD neurons generated from the KOLF2.1J line. **N** Representative trace of iPSCs evoked by 20Hz stimulation trains with zoom-in on first five pulses in KOLF2.1J. **O** Plot of cumulative iPSC amplitudes evoked by 20 Hz stimulus trains, with back-extrapolated linear fits displayed in red. **P** RRP estimate obtained from back-extrapolation of linear fits displayed in (**M**). **Q**. representative ePSC traces from PB-AD neurons induced from the GM23973 line. **R.** Representative trace of iPSCs evoked by 20Hz stimulation trains with zoom-in on first five pulses in GM23973. **S** Amplitude, (**T**) charge and (**U**) decay time constant of iPSCs in PB-AD neurons generated from the GM23973.1J line. **V** Plot of cumulative iPSC amplitudes evoked by 20 Hz stimulus trains, with back-extrapolated linear fits displayed in red. **W** RRP estimate obtained from back-extrapolation of linear fits displayed in (M). Boxplot whiskers extend the entire data range.

**Figure S3.**
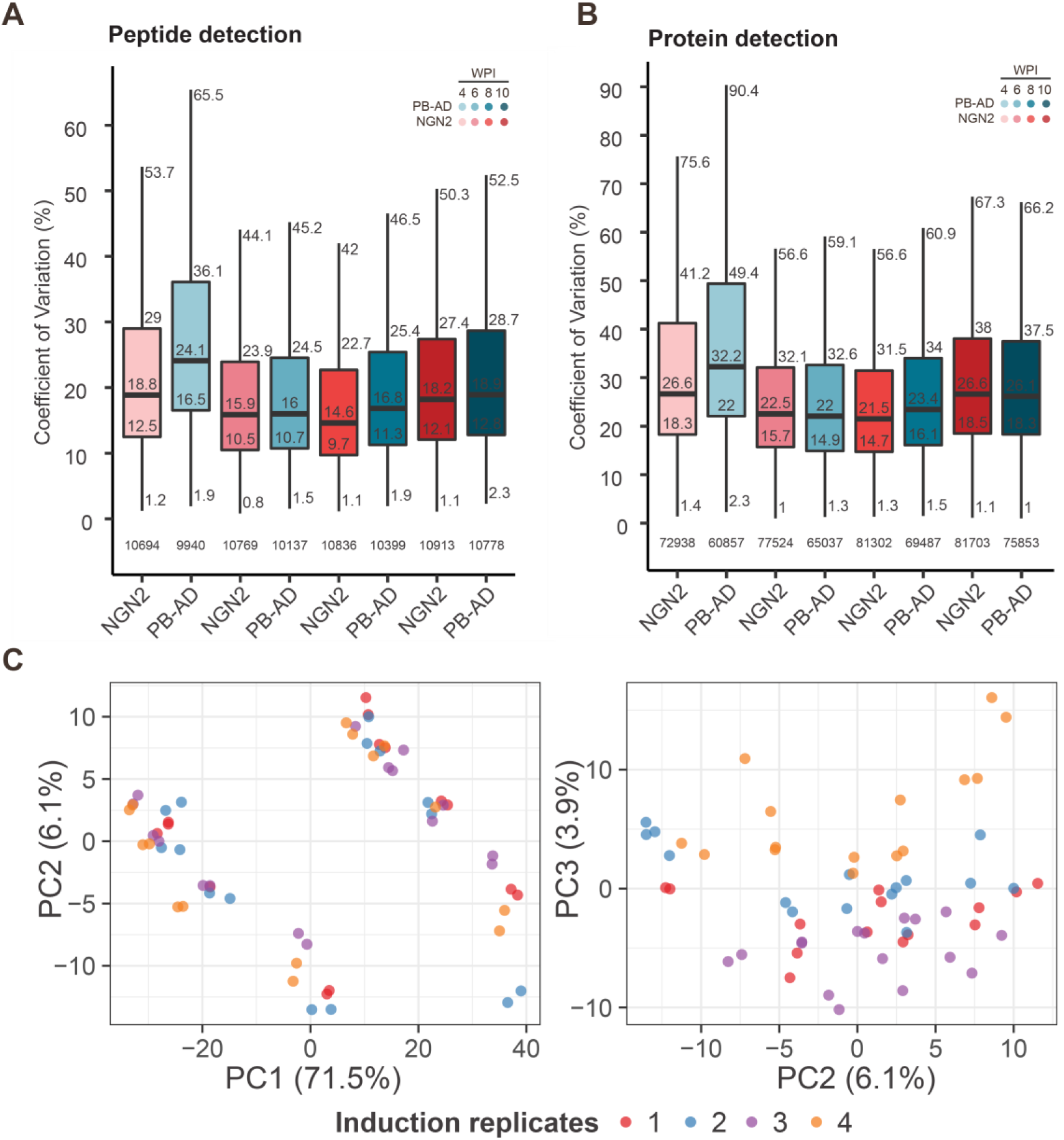
Peptide and protein detection count and variation is similar between cultures and timepoints. **A-B** Boxplots showing variation of peptide (**A**) and protein (**B**) detection for all timepoints per culture, indicating median, 1^st^ and 3^rd^ quartiles, and minimum and maximum values. Number of detected peptides and proteins are indicated in the base of the graph. Median coefficient of variation for peptide and protein detection across groups ranged from 14.6% to 32.2%, reflecting consistent detection performance across replicates. **C** PCA plots showing potential batch effects due to induction. PC1 and PC2 show even spread of different inductions but PC3 (3.9%) shows slight separation based on induction and we therefore corrected for induction effect when performing differential abundance analysis.

**Figure S4.**
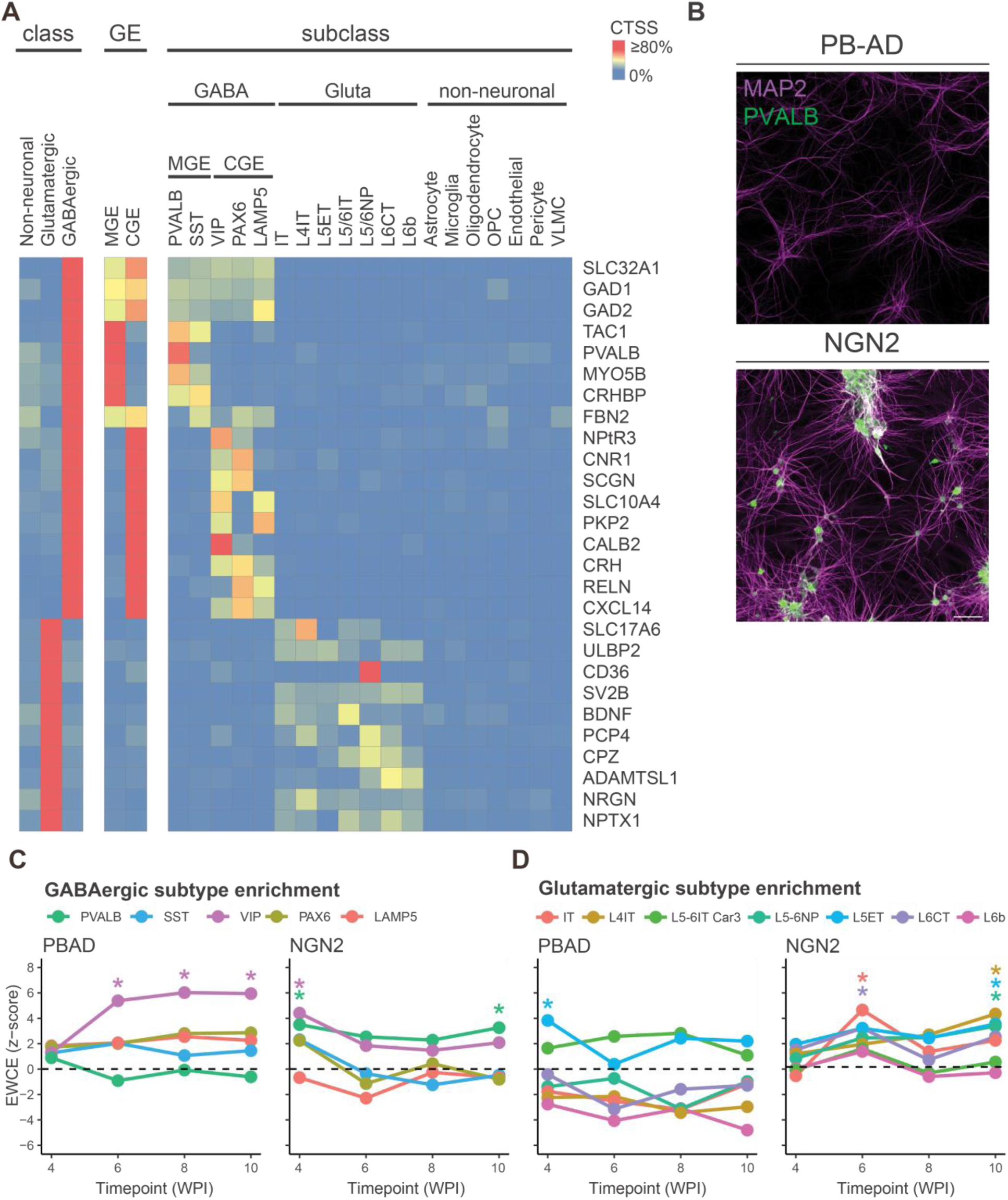
Cell type specificity scores for detected GABAergic proteins. **A**. Cell type specificity scores (CTSS) scores for detected GABAergic and glutamatergic proteins. CTSS are calculated with the expression weighted cell type enrichment tool.^41^ using single-cell RNA-sequencing data from Jorstad et al. ^40^ MGE and CGE refer to medial and caudal ganglionic eminences. **B.** Representative images of PB-AD and NGN2 neurons stained for MAP2 and PVALB. Scale bar = 100 µm. **C-D** Expression-weighted cell type enrichment (EWCE) analysis showing enrichment of proteins higher expressed in PB-AD or NGN2 for GABAergic **(C)** and glutamatergic subtypes **(D)**. PBAD-enriched proteins showed strongest enrichment for VIP^+^ interneurons at most timepoints, whereas NGN2-enriched proteins map at different timepoints to both GABAergic (PVALB^+^ and VIP^+^ interneurons) and glutamatergic subtypes. PB-AD-enriched proteins showed unexpected enrichment for L5ET neurons at 4 WPI, which reduced below significance threshold at later timepoints. IT and ET refer to intra- and extratelenchephalic-projecting neurons, and NP refers to near-projecting neurons – cell subclass annotations consistent with Jorstad et al. (2023).^40^

**Figure S5.**
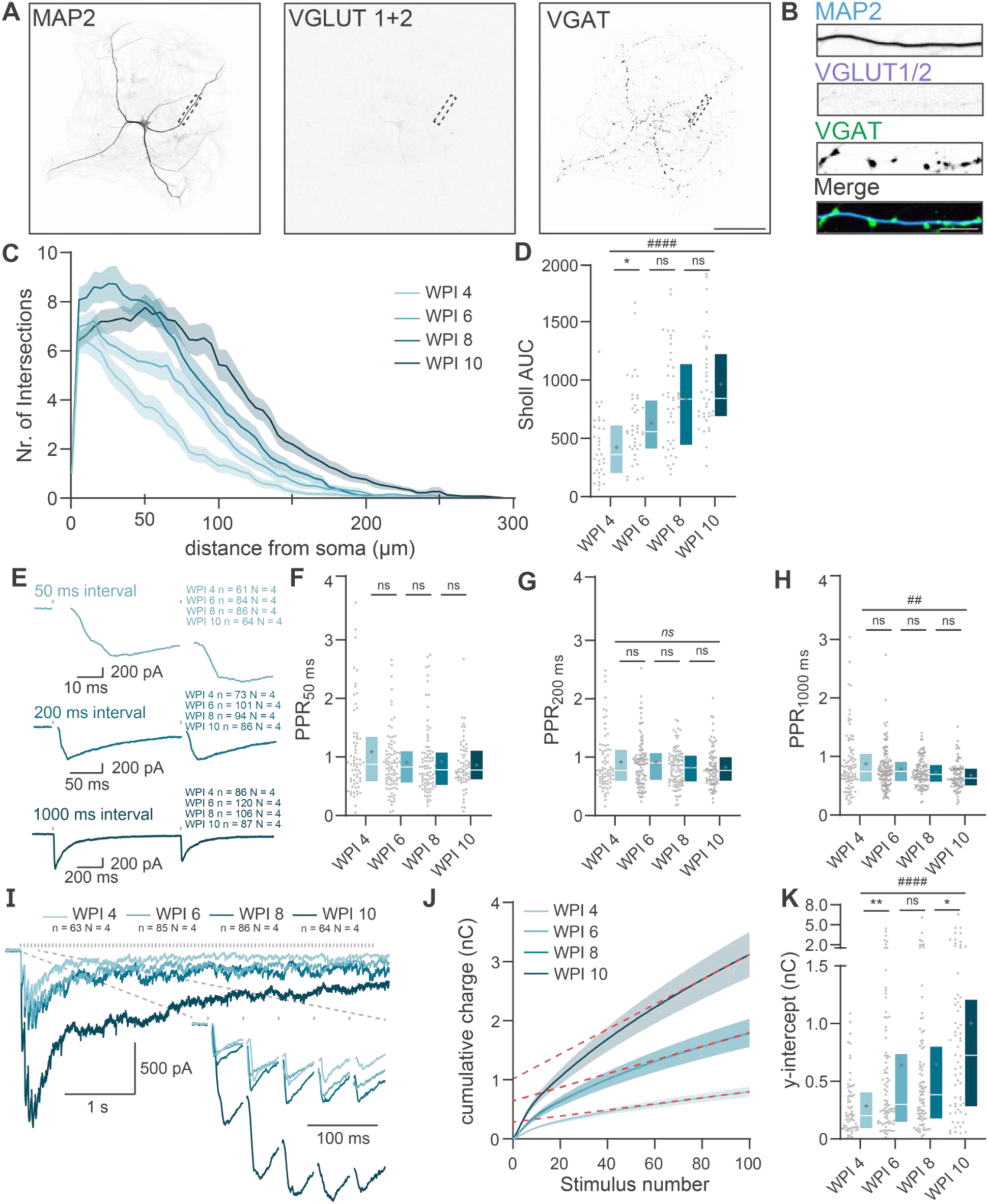
Extended morphological and functional development of PiggyBac Ascl1/Dlx2 induced neurons. **A** Representative images of an autaptic PiggyBac (PB) *Ascl1*/*Dlx2* (AD) induced neuron at week-post-induction 10, stained for MAP2 as dendritic marker, and VGAT and VGLUT1/2 as GABAergic and glutamatergic markers, respectively. Scale bar: 100 µm **B** Cut-out of dendrite marked in A to illustrate individual synaptic puncta. Scale bar: 10 µm **C** Sholl analysis of PB-AD autaptic neurons at WPI 4, 6, 8 and 10. Shaded areas represent the SEM. **D** Quantification of the area under the curve (AUC) of the Sholl histogram shown in (**C**) **E** Representative traces of paired-pulse ePSCs at a 50, 200 and 1000 ms interval at 10 WPI. **F** Quantified paired-pulse amplitude ratios (PPR) at a 50 ms, 200 ms (**G**) and 100 ms (**H**) interval at 4, 6, 8 and 10 WPI. **I** Representative traces of iPSCs evoked by 20Hz stimulus trains from PB-AD neurons at 4, 6, 8 and 10 WPI with zoom in on first five pulses. **J** Plot of cumulative iPSC charge evoked by 20 Hz stimulus trains, with back-extrapolated linear fits displayed in red. Note, WPI 4 and WPI 6 data overlap. **K** RRP estimate obtained from back-extrapolation of linear fits displayed in (**J**).

**Figure S6.**
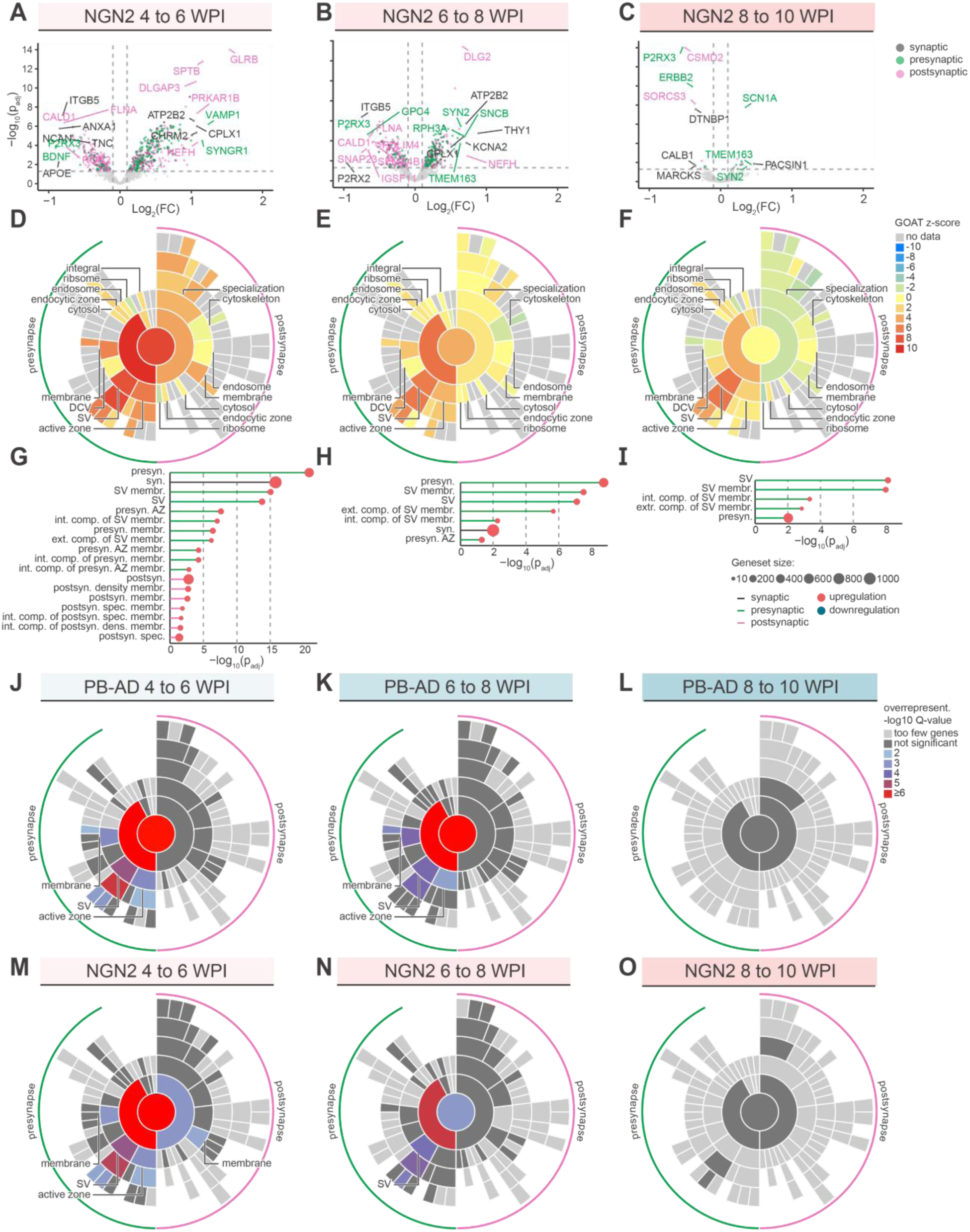
Differential expression of synaptic genes in NGN2 and PB-AD neurons. **A-C.** Volcano plots showing regulation of synaptic proteins in NGN2 cultures between 6 and 4 (**A**), 8 and 6 (**B**), and 10 and 8 WPI (**C**). Only SYNGO-annotated proteins are plotted and colour coding refer to significantly regulated synaptic (black) exclusively presynaptic (green) or postsynaptic (pink) annotations. Proteins were regarded significantly regulated when adjusted p-value < 0.05 (dashed horizontal line) and log^2^ (FC) > 0.1 or < −0.1 (dashed vertical lines). **D-F**. SYNGO plots showing enrichment scores (z-scores) derived from gene-set enrichment analysis with GOAT using SYNGO cellular compartment gene sets for 4 to 6 (**D**), 6 to 8 (**E**) and 8 to 10 WPI (**F**). Negative z-scores (blue) reflect enrichments primarily driven by downregulated proteins and vice versa for positive z-scores (red). **G-I.** Lollipop charts showing significant SYNGO terms with their corresponding adjusted p-values and the number of genes part of the gene set. **J-O.** SYNGO overrepresentation analysis of upregulated proteins in PB-AD (**J-L**) and NGN2 (**M-O**) per timepoint. Foreground are significantly upregulated proteins and background are all tested genes in corresponding contrast.

**Figure S7.**
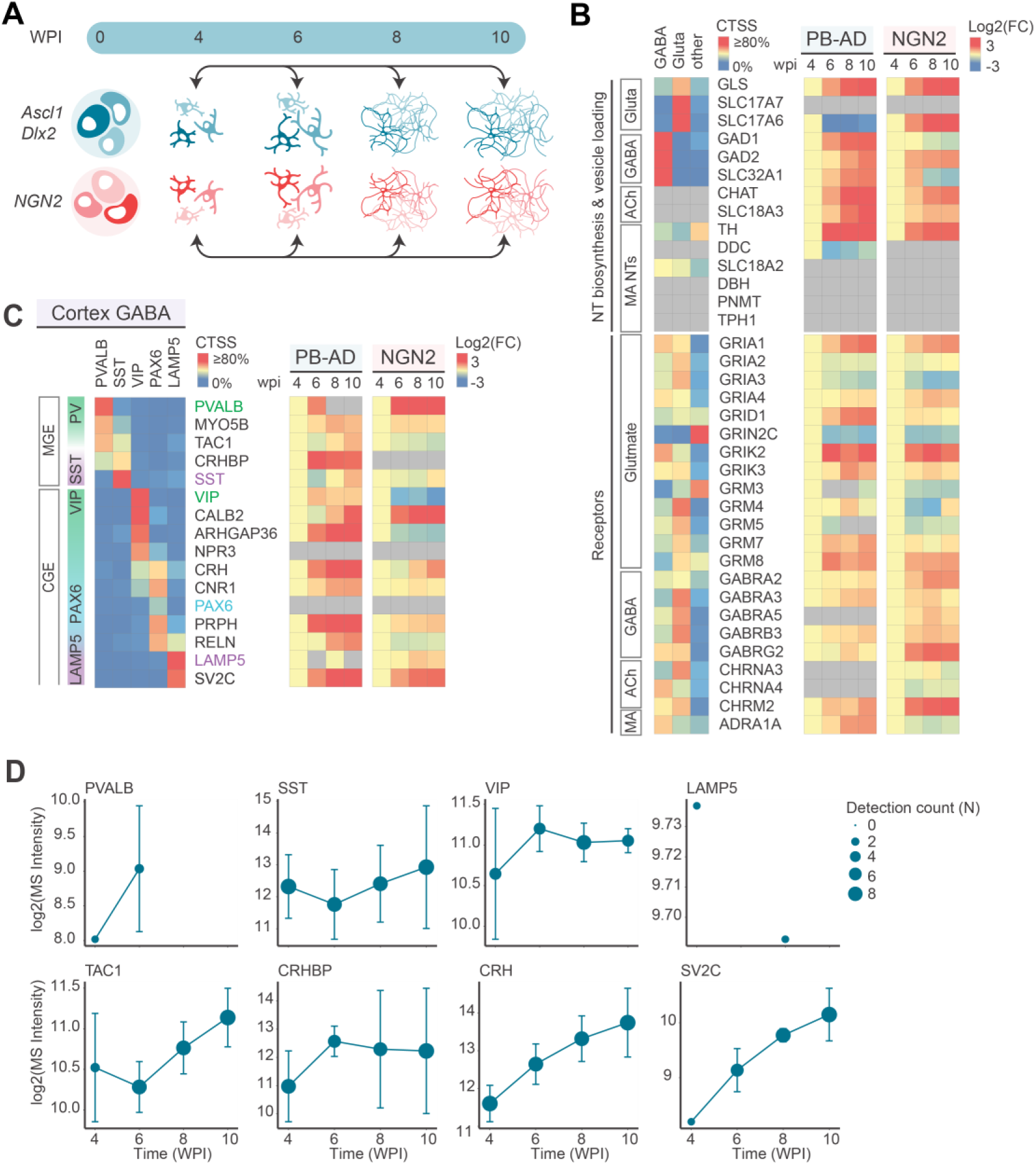
PiggyBac Ascl1/Dlx2 and NGN2 neurons express markers of diverse GABAergic neuron subtypes. **A.** Contrasts used to determine cell type specification. **B** and **C** Heatmaps that show temporal expression profiles relative to 4 WPI (log^2^ (FC)) and cell type specificity scores (CTSS). **B** Typical proteins required for neurotransmitter (NT) synthesis, vesicle loading, and receptors. Undetected proteins are visualized for NT biosynthesis and vesicle loading but not for receptors. **C** Canonical GABAergic subtype markers and proteins with high cell type specificity for GABAergic subtypes and for medial or caudal ganglionic eminences (MGE or CGE). **D.** Line plots show sum expression of canonical GABAergic subtype markers (top) and other proteins with GABAergic cell type specificity. Error bars represent SD and dot size indicate detection count.

**Supplemental table 1:** SYNGO Proteins with significant expression changes from 8 to 10 WPI in PB-AD neurons.

| Protein | new at WPI 10 | log<sub>2</sub>fc | FDR-adjusted p-value |
| --- | --- | --- | --- |
| CNTNAP4 | yes | 1.05 | 1.50E-26 |
| SLITRK1 | yes | 0.97 | 7.75E-23 |
| CDH1 | yes | 0.91 | 4.81E-20 |
| EPHA7 | yes | 0.90 | 1.26E-19 |
| ITGB3 | no | 0.80 | 1.72E-15 |
| PRRT1 | yes | 0.74 | 1.35E-13 |
| NPTX2 | no | 0.73 | 4.68E-13 |
| ITGA2 | no | 0.67 | 3.46E-11 |
| SLC17A5 | yes | 0.67 | 5.09E-11 |
| KCNQ2 | yes | 0.65 | 1.77E-10 |
| ELFN1 | no | 0.65 | 2.09E-10 |
| CYTH1 | no | 0.65 | 2.21E-10 |
| TNC | no | 0.64 | 3.74E-10 |
| NRGN | yes | 0.63 | 6.71E-10 |
| NGDN | yes | 0.63 | 6.73E-10 |
| GPR158 | no | 0.62 | 1.43E-09 |
| RELN | yes | 0.57 | 3.53E-08 |
| SCG2 | no | 0.78 | 2.18E-03 |
| TMEM163 | no | 0.62 | 7.64E-03 |
| PTPRN | no | 0.59 | 2.24E-02 |
| NEFH | no | 0.40 | 2.24E-02 |
| IGSF8 | no | 0.40 | 2.24E-02 |
| CNTNAP1 | no | 0.34 | 3.78E-02 |
| ATP2B4 | no | 0.30 | 3.78E-02 |

